# Genomic evolution of the globally disseminated multidrug-resistant *Klebsiella pneumoniae* clonal group 147

**DOI:** 10.1101/2021.07.03.450759

**Authors:** Carla Rodrigues, Siddhi Desai, Virginie Passet, Devarshi Gajjar, Sylvain Brisse

## Abstract

**Background:** The rapid emergence of multidrug-resistant *Klebsiella pneumoniae* (Kp) is largely driven by the spread of specific clonal groups (CG). Of these, CG147 includes 7-gene MLST sequence types ST147, ST273 and ST392. CG147 has caused nosocomial outbreaks across the world, but its global population dynamics remain unknown. Here, we report a pandrug-resistant ST147 clinical isolate from India (strain DJ) and define the evolution and global emergence of CG147.

**Methods:** Antimicrobial susceptibility testing (EUCAST guidelines) and genome sequencing (Illumina and Oxford Nanopore technologies, Unicycler assembly) were performed on strain DJ. Additionally, we collated 217 publicly available CG147 genomes (NCBI, May 2019). CG147 evolution was inferred within a temporal phylogenetic framework (BEAST) based on a recombination-free sequence alignment (Roary/Gubbins). Comparative genomic analyses focused on resistance and virulence genes and other genetic elements (BIGSdb, Kleborate, PlasmidFinder, PHASTER, ICEFinder and CRISPRCasFinder).

**Results:** Strain DJ had a pandrug resistance phenotype. Its genome comprised 7 plasmids and 1 linear phage-plasmid. Four carbapenemase genes were detected: *bla*_NDM-5_ and 2 copies of *bla*_OXA-181_ in the chromosome, and a second copy of *bla*_NDM-5_ on an 84 kb IncFII plasmid. CG147 genomes carried a mean of 13 acquired resistance genes or mutations; 63% carried a carbapenemase gene and 83% harbored *bla*_CTX-M_. All CG147 genomes presented GyrA and ParC mutations and a common subtype IV-E CRISPR-Cas system. ST392 and ST273 emerged in 2005 and 1995, respectively. ST147, the most represented phylogenetic branch, was itself divided into two main clades with distinct capsular loci: KL64 (74%, DJ included, emerged in 1994 and disseminated worldwide, with carbapenemases varying among world regions) and KL10 (20%, 2002, predominantly found in Asian countries, associated with carbapenemases NDM and OXA-48-like). Further, subclades within ST147-KL64 differed in the yersiniabactin locus, OmpK35/K36 mutations, plasmid replicons and prophages. The absence of IncF plasmids in some subclades was associated with a possible activity of a CRISPR-Cas system.

**Conclusions:** *K. pneumoniae* clonal group CG147 comprises pandrug- or extensively-resistant isolates and carries multiple and diverse resistance genes and mobile genetic elements, including chromosomal *bla*_NDM-5_. Its emergence is driven by the spread of several phylogenetic clades marked by their own genomic features and specific temporo-spatial dynamics. These findings highlight the need for precision surveillance strategies to limit the spread of particularly concerning CG147 subsets.

## INTRODUCTION

The increasing number of antimicrobial resistant infections by *Klebsiella pneumoniae* (Kp), especially by extended-spectrum beta-lactamases (ESBL)- and carbapenemase-producing Kp, led to the declaration of Kp as an ‘urgent threat’ and ‘priority pathogen’ by public health agencies (CDC, 2013; WHO, 2014). Molecular analyses of Kp isolates has evidenced that the rapid emergence of multidrug resistant (MDR) Kp is largely driven by the geographic spread of successful clonal groups (CG; e.g., CG15, CG101, CG147, CG258, CG307) (Wyres et al., 2020), some of them carrying epidemic resistance plasmids (Navon-Venezia et al., 2017). In order to treat MDR Kp infections, last-resort drugs such as polymyxins (especially colistin) and tigecycline are used (Poirel et al., 2017; Wyres et al., 2020). Consequently, resistance is also observed to these last resort drugs, especially to colistin, and may culminate in the emergence and spread of pandrug-resistant strains (Magiorakos et al., 2012; Poirel et al., 2017). Pandrug-resistant Kp strains leave few or no therapeutic options and are associated with high mortality rates (Avgoulea et al., 2018; de Man et al., 2018; Guducuoglu et al., 2018; Lázaro-Perona et al., 2018; Zowawi et al., 2015).

The 7-gene MLST sequence type (ST) ST147 has been recognized as a globally distributed antimicrobial resistance clone (Peirano et al., 2020), and is closely related to ST273 and ST392, which themselves comprise MDR isolates. Based on genomic classifications, these three STs are grouped into the CG147 clonal group (Bialek-Davenet et al., 2014; Wyres, Wick, et al., 2019). The earliest reports of CG147 isolation date from 2008-2010 in Hungary, and correspond to ciprofloxacin-resistant CTX-M-15-producing ST147 isolates, which had been disseminating since 2005 (Damjanova et al., 2008; Szilágyi et al., 2010). Between 2010 and 2014, CG147 (mainly ST147) was described worldwide in association with several carbapenemases (Munoz-Price et al., 2013). Most reported CG147 isolates are from clinical samples, although some were found in companion animals, river waters, chimpanzees, poultry animals and poultry environment (Baron et al., 2021; Ovejero et al., 2017; Sato et al., 2018; Suzuki et al., 2020; Zhang et al., 2019; Zogg et al., 2018; Zurfluh et al., 2015). The above studies were locally restricted and so far, no study of the global spread and genome dynamics of this clone was performed.

Here, we report a pandrug-resistant clinical isolate from India belonging to ST147, and investigate the genomic evolution and antimicrobial resistance gene dynamics in the global CG147 population. We also analyze the phylogenetic context of virulence-associated genomic features, CRIPSR loci and mobile genetic elements in this successful Kp sublineage.

## MATERIAL AND METHODS

### Isolation and phenotypic and genomic characterization of strain DJ

Isolate DJ was recovered from the urine of a 45-years old female patient diagnosed with a urinary tract infection (UTI) in Vadodara (Gujarat, India) in October 2016. The isolate was confirmed to be Kp by biochemical tests and 16S *rRNA* sequencing. Antibiotic susceptibility tests performed using a semi-automated commercial system (Vitek, BioMérieux) revealed resistance to all antibiotics tested. Confirmatory antimicrobial susceptibility tests were carried out for colistin and tigecycline using broth dilution; and for fosfomycin (agar supplemented with 25 mg/L glucose 6-phosphate sodium salt) using agar dilution method. Disk diffusion method was used for the other antimicrobial classes (penicillins, cephalosporins, carbapenems, monobactams, fluoroquinolones, aminoglycosides, macrolides, tetracyclines, phenicols and inhibitors of the folic acid pathway). Results were interpreted using both the Clinical and Laboratory Standards Institute (CLSI, 2016) and the European Committee on Antimicrobial Susceptibility Testing (2018) (http://www.eucast.org/) guidelines.

DNA extraction was performed using XpressDNA Bacteria kit (MagGenome Technologies Pvt Ltd., India). Whole-genome sequencing data were generated using (i) an Illumina NextSeq-500 platform with a 2 × 150 nt paired-end protocol (Nextera XT library, Illumina, San Diego, CA); and (ii) long-read Oxford Nanopore sequencing using MinION device integrated with a FLO-MIN-106 flow cell and libraries prepared using a 1D ligation sequencing kit (SQK-LSK109) following the protocol for 1D genomic DNA (gDNA) long reads without BluePippin (Oxford Nanopore Technologies, New York, NY, USA). *De novo* assemblies of the reads were obtained using SPAdes v3.12.0 (Prjibelski et al., 2020) for Illumina data and using Unicycler v0.4.4 (Wick et al., 2017) for hybrid assembly. Assembled sequences were annotated using Prokka v1.12 (Seemann, 2014). Reads and assembly were deposited at the European Nucleotide Archive database (under the BioProject PRJEB41234).

### Global dataset of publicly-available genomic sequences

All publicly available CG147 Kp genomes from NCBI RefSeq repository of genome assemblies (May 2019) were downloaded. From the 245 CG147 genomes available, duplicated (n=5) and poor-quality genomes (n=6; genome size and G+C content not matching with Kp and/or more than >1000 contigs), and those without attached isolation year (n=17) were excluded. The final dataset comprised 218 genomes, including strain DJ. Sample information, accession numbers, and biological characteristics of the genomes are given in **Table S1**.

### Phylogenetic analyses

For phylogenetic analyses, a core-genome alignment based on the concatenation of 4,529 core genes was obtained using Roary v3.12 (Page et al., 2015) using a blastP identity cut-off of 90% and core genes defined as those being present in more than 90% of the isolates. Recombination events were removed from the core-genome alignment using Gubbins v2.2.0 (Croucher et al., 2015). The final recombination-free alignment comprised 8,450 single-nucleotide variants (SNVs) and was used to construct a maximum-likelihood phylogenetic tree using IQ-TREE v1.6.11 (model GTR+F+ASC+G4). The tree was rooted with a Kp ST258 NJST258_2 (accession number: GCF_000597905.1) and a Kp ST37 INF042 (GCF_002752995.1) (**Figure S1**).

To evaluate the strength of the temporal signal of our molecular phylogeny, we first conducted a linear regression analysis of the root-to-tip genetic distances as a function of the sample collection year, using TempEst v1.5.3 (http://tree.bio.ed.ac.uk/software/tempest/) (**Figure S2**). The final recombination-free alignment was then subjected to Bayesian phylogenetic analysis using BEAST v2.6.1 (run with a Markov chain Monte Carlo length of 1× 10^9^, sampling every 5×10^3^ steps) (Bouckaert et al., 2019). We used model parameters that had the best fit: GTR substitution model, lognormal relaxed clock and constant population size. Parameters estimates were computed using Tracer v1.7.1, and a maximum clade credibility tree was obtained with TreeAnnotator v2.6.0. and visualized in FigTree v1.4.4.

### MLST and genomic analyses of resistance, virulence and other genetic elements

Multilocus sequence typing (MLST, 7 genes) was performed using the Institut Pasteur *Klebsiella* MLST (Diancourt et al., 2005) database (https://bigsdb.pasteur.fr/klebsiella/). Kleborate (Lam et al., 2020) and BIGSdb analytical tools (https://bigsdb.pasteur.fr/klebsiella/) (Jolley et al., 2018) were used to define the presence of antimicrobial resistance, heavy metal tolerance, virulence genes and to characterize the capsular gene cluster. Geneious Prime 2019.1.1 software (https://www.geneious.com) was used for further manual curation of antibiotic resistance genes, and ISFinder (https://isfinder.biotoul.fr) was used to look for the insertion sequences in the resistance genes or in their genetic context. Plasmid replicons were detected using PlasmidFinder (https://cge.cbs.dtu.dk/services/PlasmidFinder/) (Carattoli et al., 2014), whereas prophages, integrative and conjugative elements (ICEs) and CRISPRs were identified using PHASTER (https://phaster.ca) (Arndt et al., 2016), ICEfinder (https://db-mml.sjtu.edu.cn/ICEfinder/ICEfinder.html) and CRISPRCasFinder (https://crisprcas.i2bc.paris-saclay.fr/CrisprCasFinder/Index) (Couvin et al., 2018), respectively. To depict co-resistance genotypes and plasmid networks, we constructed a correlation matrix for binary variables (1, presence; 0, absence) using the ‘corr.test’ function (Pearson method, which for a pair of binary variables compares to the Phi coefficient) from the ‘corrplot’ R package. Significant correlations were visualized with the ‘corrplot’ function from the same package. Statistical analyses to check the association of the different categorical variables within the phylogeny groups were calculated using the *χ*^2^ test (*P* values of <0.05 were considered statistically significant).

## RESULTS

### Phenotypic and genomic features of pan-drug resistant strain DJ

Strain DJ was resistant to all tested antimicrobial agents, including last-resort antimicrobials such as carbapenems, colistin, tigecycline and fosfomycin (**Table 1**), and is therefore pandrug-resistant (Magiorakos et al., 2012). To define its genetic mechanisms of resistance, a hybrid complete genome assembly was produced (**Figure S3a**). The 5.7 Mb sequence was 56.9% G+C-rich and made up of one chromosome and 6 circularized plasmids [123kb IncFII(pKPX1); 57 kb IncR; 5.6 kb ColRNAI; 4.7 kb ColRNAI; 2.0 kb ColpVC; and 1.5 kb ColMG828]. In addition, there were two non-circularized contigs: a 84 kb IncFII plasmid and a 57 kb contig corresponding to a N15-*like* phage-plasmid (P-P) coding for a protelomerase (*telN*) responsible for the maintenance of its linear genome (Pfeifer et al., 2021). PHASTER identified four other prophages within the chromosome. Phylogenetic analysis of strain DJ showed it belonged to *K. pneumoniae sensu stricto* (*i.e.,* phylogroup Kp1) and to the ST147-KL64 lineage previously described as endemic in India (Peirano et al., 2020).

**Table 1.** Antimicrobial susceptibility of *Klebsiella pneumoniae* strain DJ, and genes potentially conferring resistance.

| Class and antimicrobial agent | Diameter (mm) or MIC (µg/ml) | Interpretation<sup>1</sup> | Associated resistance genes (copy number) |
| --- | --- | --- | --- |
| **Beta-lactams** |  |  | *bla*<sub>NDM-5</sub> (2), *bla*<sub>OXA-181</sub> (2), *bla*<sub>CTX-M-15</sub> (3), *bla*<sub>TEM-1</sub> (2), *bla*<sub>SHV-11</sub> (1), disrupted *ompK35*, mutation in *ompK36* |
| Ampicillin | 9 | R |  |
| Piperacillin | 13 | R |  |
| Amoxycillin-clavulanic acid | 11 | R |  |
| Ticarcillin-clavulanic acid | 14 | R |  |
| Piperacillin-tazobactam | 14 | R |  |
| Cefuroxime | 11 | R |  |
| Cefotaxime | 12 | R |  |
| Ceftazidime | 15 | R |  |
| Cefepime | 15 | R |  |
| Aztreonam | 14 | R |  |
| Ertapenem | 12 | R |  |
| Imipenem | 14 | R |  |
| Meropenem | 15 | R |  |
| **Aminoglycosides** |  |  | *rmtB* (1), *rmtF* (2), *aac(6')-Ib* (2), *aadA2* (3), *strAB*(2) |
| Amikacin | 10 | R |  |
| Gentamycin | 10 | R |  |
| Tobramycin | 9 | R |  |
| **Quinolones and Fluoroquinolones** |  |  | *gyrA* and *parC* mutations |
| Nalidixic acid | 11 | R |  |
| Ofloxacin | 11 | R |  |
| **Folate pathway inhibitors** |  |  | *sul1* (3), *sul2* (1), *dfrA12* (3) |
| Trimethoprim/Sulfamethoxazole | 19 | R |  |
| Trimethoprim | 8 | R |  |
| **Polymyxins** |  |  | disrupted *mgrB* |
| Colistin | MIC=4 | R |  |
| **Tetracyclines** |  |  | *ramR* mutation<sup>2</sup> |
| Tigecycline | MIC=16 | R |  |
| **Phenicals** |  |  | *catA2* (1), *catB* (2) |
| Chloramphenicol | 12 | R |  |
| **Other** |  |  |  |
| Fosfomycin | MIC=128 | R | *fosA* |
| Rifampicin | NI | - | *arr-2* (2) |
| Macrolides | NI | - | *mphA* (1), *ermB* (1) |

NI, not included in the antimicrobial susceptibility testing panel; R, resistant; <sup>1</sup>Interpretations were based on CLSI and EUCAST guidelines; <sup>2</sup> Nucleotide mutation resulting in a premature stop codon.

Strain DJ harbored four carbapenemase genes, corresponding to two copies of each *bla*_NDM-5_ and *bla*_OXA-181_. Additionally, three copies of *bla*_CTX-M-15_ were detected (**Table 1**). The above genes were localized as follows (**Figure S3, Figure 1**): (i) *bla*_NDM-5_: one copy in the chromosome, and the second copy in the 84 kb IncFII plasmid; (ii) *bla*_OXA-181_: two copies in the chromosome; and (iii) *bla*_CTX-M-15_: two copies in the chromosome and one copy in the 57 kb IncR plasmid. In addition, a 61 kb region from the IncFII plasmid, comprising several antimicrobial resistance genes included in a class 1 integron and the replication machinery of the plasmid, was duplicated in the chromosome (**FigureS3a,c**).

**Figure 1.**
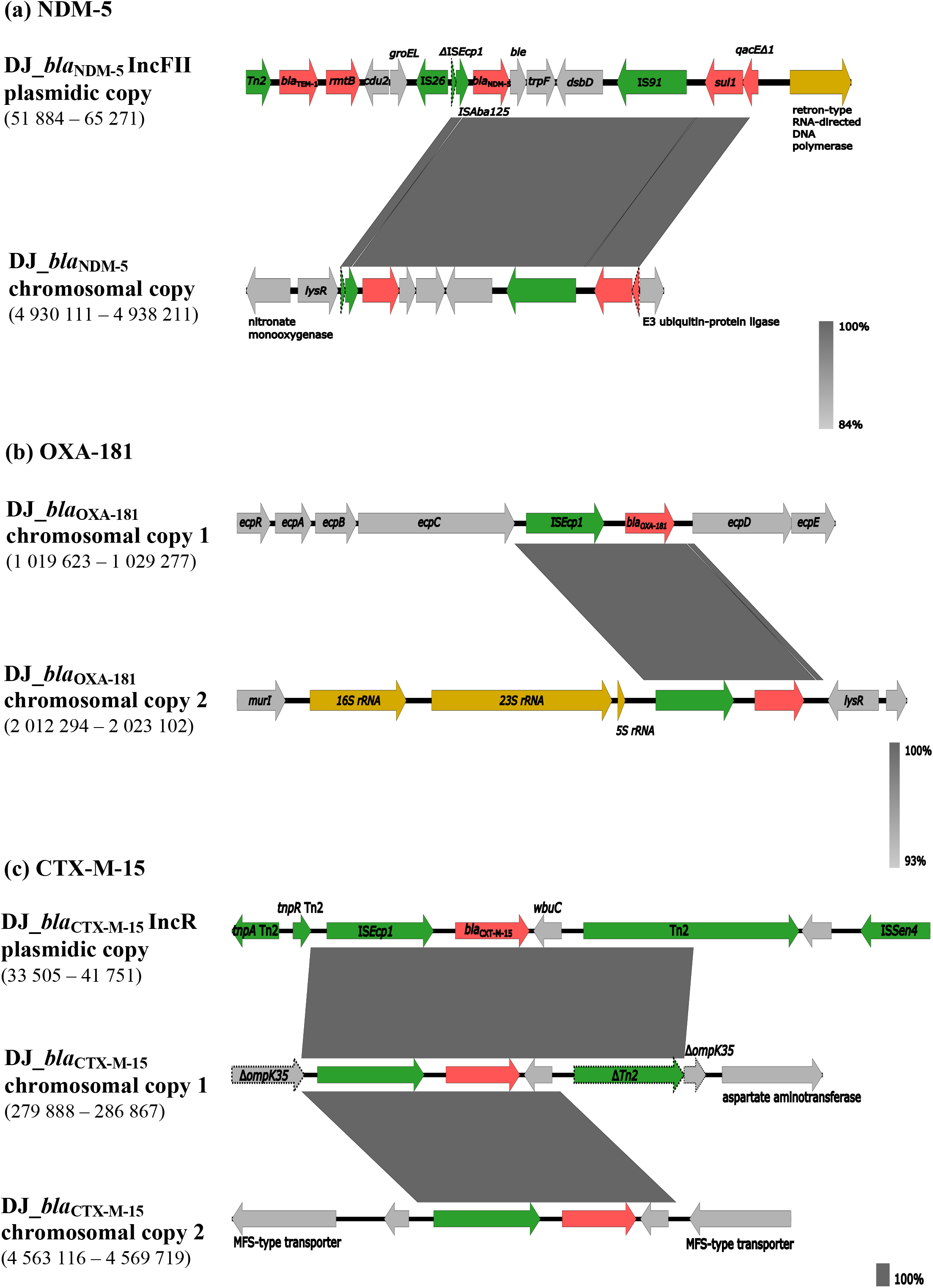
Genetic context of the different copies of *bla*_NDM-5_ **(a)**, *bla*_OXA-181_ **(b)** and *bla*_CTX-M-15_ **(c)** genes identified in *K. pneumoniae* strain DJ. Within each panel, the chromosomal and/or plasmid-encoded copies of *bla*_NDM-5_, *bla*_OXA-181_ and *bla*_CTX-M-15_ detected in DJ are compared between them. Predicted open reading frames (ORFs) are represented on each line by colored arrows, with arrowheads indicating the direction of transcription: antimicrobial resistance genes (red), mobile genetic elements (MGEs) or gene mobilization-related genes (green); RNA-binding proteins (dark yellow); other functions (light gray). Disrupted genes are outlined with dotted lines. Dark gray blocks connecting the distinct gene regions represent homology levels, as indicated in the gradient key. The nucleotide positions of the represented regions are indicated at the left below the copy descriptors. Figures were created using the Easyfig (https://mjsull.github.io/Easyfig/).

Additional molecular determinants of antimicrobial resistance were observed in strain DJ. Most notably, the gene *mgrB* was disrupted by an IS*5* transposase (1057 bp), consistent with the observed colistin resistance phenotype (MIC=4 μg/ml). This strain also carried *rmtF, rmtB, strA, strB* and *aadA2* genes, associated with resistance to aminoglycosides including amikacin **(Table 1)**. Quinolone resistance determining region (QRDR) mutations were observed, leading to GyrA-83I and ParC-80I amino acid alterations. Last, there was a premature stop codon caused by a A580T substitution in RamR, a negative regulator of RamA, itself a transcriptional activator of the *acrAB* genes. Higher production of AcrAB increases the efflux of tigecycline (Villa et al., 2014), consistent with the resistance phenotype observed for this agent (MIC=16 μg/ml).

Regarding virulence genes, strain DJ harbored a complete yersiniabactin gene cluster (*ybt*10) located on an ICE*Kp*4 mobile genetic element (**Figure S3a**). Type 1 (*fimAICDFGH*) and type 3 (*mrkABCD*) fimbriae gene clusters were also observed, but none of the *rmpACD*, aerobactin and salmochelin cluster genes, typically associated with hypervirulence, were present.

### Time-scaled phylogenetic structure of the global population of sublineage CG147

We investigated the evolutionary origins of strain DJ within the global diversity of CG147 using 217 publicly available genomes of isolates collected between 2002 and 2018. CG147 genomes were mainly isolated from human samples (90%, 196/218) and hospital environment (6%, 12/218), mostly in European (40%), Southeastern Asia (17%), Southern Asia (11%) and Northern America (10%) countries.

CG147 was deeply structured into three main branches (**Figure 2**), each corresponding to a single MLST sequence type: ST147, ST273 and ST392; the two latter are *tonB* variants of ST147. The number of genome-wide nucleotide substitutions was associated with isolation dates (root-to-tip regression analysis: *R*^2^ =0.1123, **Figure S2**), enabling to infer a time-scaled phylogeny (**Figure 3**). The evolutionary rate within CG147 was estimated at 1.45×10^−6^ substitutions/site/year (95% HPD, 1.12×10^−6^ – 1.78 ×10^−6^), corresponding to 6.2 SNPs per genome per year. The last common ancestor of CG147 was estimated around year 1896, with a large uncertainty (95% highest posterior density (HPD): 1817-1962). The ST273 lineage was the first to diverge, whereas ST147 and ST392 shared a common ancestor, estimated around 1921 (95% HPD, 1868-1970) (**Figure 3**).

**Figure 2.**
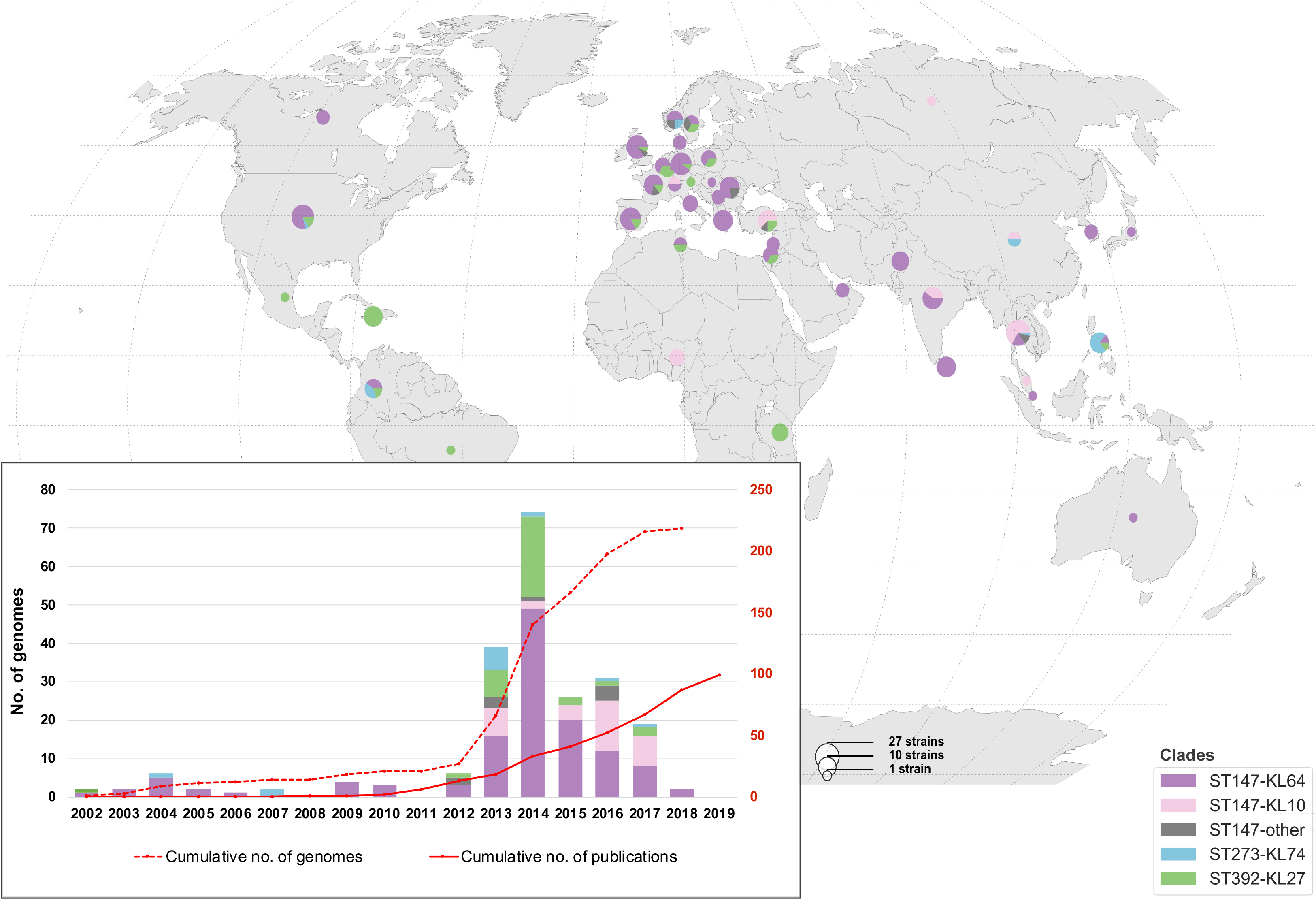
Geographic **(main)** and temporal **(inset)** distribution of genomes included in this study. The pie charts represent the frequency of each CG147 clade in each country (see size and color keys). **Inset:** Bars represent the number of isolates per year for which genome assemblies were available (NCBI RefSeq) as of May 2019, coloured by clade. Red lines represent (solid line) the number of PubMed-indexed records as of March 2020 (identified using the search criteria ‘*Klebsiella pneumoniae*’ and ‘ST147’ or ‘ST392’ or ‘ST273’, resulting in a total of 99 distinct entries); and (dotted line) the cumulative number of genomes. The scale on the left Y-axis refers to total number whereas the one on the right refers to cumulative numbers.

**Figure 3.**
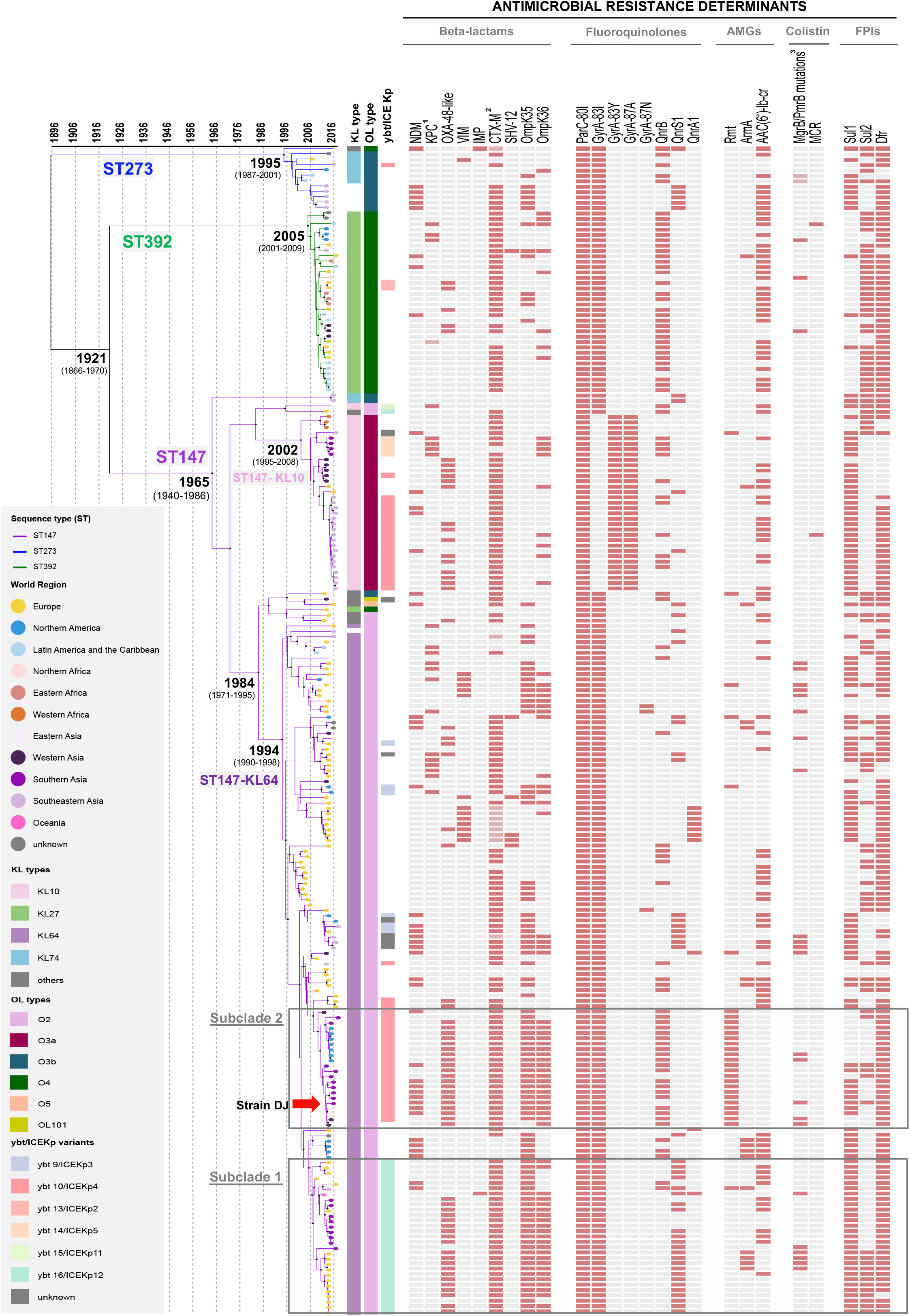
Time-scale phylogeny of 218 CG147 genomes and their epidemiological and molecular characteristics. The phylogeny was obtained using the BEAST tool. The three main branches correspond to the three main sequence types (ST). Tree tips are colored by world region of isolation (see key). Black dots on main nodes indicate ≥95% posterior probability. The grey boxes delineate subclades 1 and 2, as indicated. Capsular (KL) and O-antigen (O) locus types and the yersiniabactin-carrying ICE*Kp* elements are colored according to their variants as shown in the legend. Antimicrobial resistance determinants are indicated by colored rectangles when present. In the antimicrobial resistance determinants columns:^1^ dark pink indicates *bla*_KPC-2_ and light pink indicates *bla*_KPC-3;_ ^2^ dark pink indicates *bla*_CTX-M-15_ and dark pink indicates other *bla*_CTX-M_ variants; ^3^dark pink indicates *mgrB* mutations and dark pink indicates *pmrB* mutations. AMGs, aminoglycosides; FPIs, folate pathway inhibitors.

ST147 was the most represented (79%, 172/218 genomes) and geographically widespread lineage (**Figure 2**), and emerged around 1965 (95% HPD, 1940-1986). The phylogenetic structure within ST147 revealed five main clades characterized by distinct capsular (KL type) and liposaccharide O antigen loci. Clades KL64-O2 (74%; 128/172) and KL10-O3a (20%, 34/172) emerged in recent years: 1994 [95% HPD, 1990-1998] and 2002 (95% HPD, 1995-2008), respectively. Whereas KL64-O2 genomes were predominantly from Europe (54%), KL10-O3a was mainly sampled from Asia (85%; **Figure 2**).

The ST392 and ST273 branches emerged in 2005 (95% HPD, 2001-2009) and 1995 (95% HPD, 1987-2001), respectively. ST392 (16% of genomes, 34/218) is distributed globally and harbors a KL27 capsular gene cluster and a O4 antigen. In contrast, ST273 (6%, 12/218) was predominantly found in Asia (64%) and carries KL74 and O3b gene clusters. Of note, a group of closely related genomes from the Philippines (n=5) lacked a capsular gene cluster, with only *ugd* being detected. This gene had 100% identity with the *ugd* gene from the KL74 reference strain, suggesting a recent loss of the capsular gene cluster.

### Acquired antimicrobial resistance genes and their evolutionary dynamics within CG147

All genomes presented QRDR alterations in GyrA and ParC. The topoisomerase ParC 80I alteration was fully conserved, whereas the GyrA gyrase subunit 83I amino-acid change was observed in all genomes except for clade ST147-KL10, which had 83Y, caused by an ATC to TAC codon change. In addition, this clade had an 87A alteration (**Figure 3**).

The number of acquired antimicrobial resistance genes or mutations among CG147 genomes ranged from 2 to 23 (mean: 13; **Table S1**). Regarding beta-lactam resistance, in addition to the conserved chromosomal *bl*a_SHV-11_ gene, a majority of CG147 isolates carried a *bla*_CTX-M_ (n=181, 83%), 94% of which was the *bla*_CTX-M-15_ variant. In addition, 63% (n=137) genomes harbored at least one carbapenemase gene, with 14% (n=19) of these harboring more than one copy of the same carbapenemase gene, and/or two or more carbapenemase genes from different families (**Table S1**). Noteworthy, carbapenemase genes were significantly more frequent in ST147 (69%) compared to ST392 (35%) and ST273 (58%; *p=*0.0005). The two main ST147 clades were similar to this respect (ST147-KL10: 74%; ST147-KL64: 71%; *p=*0.78). However, their carbapenemase genes were distinct across world regions: there was a predominance of *bla*_NDM_ in Southeastern Asia and Northern America, whereas the combination of *bla*_NDM_ and *bla*_OXA-48-like_ was almost exclusively detected in Southeastern Asia, as observed in strain DJ. In contrast in Europe, *bla*_OXA-48-like,_ *bla*_KPC-2_ and *bla*_VIM-1/-27_ were the most frequent carbapenemases (**Figure S4; Table S1**).

Different y*bt*/ICE*Kp* subtypes (associated with hypervirulence) and OmpK35/K36 mutations (associated with multidrug resistance) were observed within the ST147-KL64 clade (**Figure 3; Table S1**). First, a group of 29 genomes (denominated subclade 1; mean number of SNPs among them: 57), emerged around 2007, was characterized by the presence of *ybt*16/ICE*Kp*12 and an altered OmpK35 protein, due to a deletion of 2 nt resulting in a premature stop codon. Within this subclade itself, a subgroup of genomes (n=19/29) harbored *bla*_OXA-48-like_ genes and the OmpK36GD mutation observed previously (Fajardo-Lubián et al., 2019). Second, a group of 22 genomes emerged around 2009 (subclade 2; mean of 60 SNPs) was defined by the presence of *ybt*10/ICE*Kp*4 and an OmpK35 gene disrupted by IS*Ecp1*-*bla*_CTX-M-15_ (**Figure 1, Figure 3, Table S1**). As observed in subclade 1, all subclade 2 isolates carrying *bla*_OXA-181_ shared the same OmpK36TD mutation (Fajardo-Lubián et al., 2019). Subclades 1 and 2 also differed in plasmid replicon content: whereas the former were rich in IncR (90%) and IncHIB/IncFIB(Mar) (41%), in contrast IncFII (pKPX1) (82%), IncFII (59%) and IncR (64%) were frequent in the latter (**Table S1**). Differently, the remaining ST147-KL64 genomes (n=77) often carried IncFIB_K_ (65%), IncFII_K_ (49%; a common pKPN-3-derived plasmid found in Kp harboring *pco* and *sil* clusters; (Navon-Venezia et al., 2017; Rodrigues et al., 2014) and IncFIA(HI1) (38%).

Strain DJ belonged to subclade 2 and was phylogenetically closely related (<28 SNPs) to five other isolates recovered between 2014 and 2015 in different Asian countries (**Figure S3b; Figure3**) and described as extremely drug resistant or pandrug-resistant (Alfaresi, 2018; Nahid et al., 2017; Zowawi et al., 2015), with identical plasmids being observed among them but no chromosomal integration of *bla*_NDM-5_ (**Figure S3**).

### Convergence of antimicrobial resistance and virulence

Whereas the yersiniabactin virulence factor gene cluster was rare amongst genomes of ST392 (6%; 2/34) and ST273 (8%; 1/12), it was observed in 53% of ST147 genomes. There were two predominant variants (*ybt*16/ICE*Kp*12 and *ybt*10/ICE*Kp*4 associated with subclades 1 and 2, respectively), and four minority ones (**Figure 3; Table S1**).

Two isolates with hypervirulence genotypes, defined by the presence of *rmpA* and/or *rmpA2* or aerobactin, were observed: KpvST147L (ST147-KL14, 2016, United Kingdom) and B-8658 (ST147-KL10, 2014, Russia). The plasmid from KpvST147L was fully sequenced previously (GenBank accession number NZ_CM007852) and carries *rmpA*, *rmpA2* and aerobactin; it corresponds to a 343 kb IncFIB-IncHIB (pNDM-MAR) plasmid (**Figure S5**) (J. F. Turton et al., 2018). This plasmid was recently described in three ST147 *Kp* isolates recovered in 2018-2019 in the United Kingdom and in other sublineages (J. Turton et al., 2019). Here, we found that Russian strain B-8658 (ST147-KL10, 2014) also acquired the pNDM-MAR plasmid, but it lacked the *rmpA* gene (**Figure S5**).

### Co-occurrence of antimicrobial resistance and heavy metal tolerance genes and plasmids

Antimicrobial resistance genes, mutations and plasmids co-occurred in a structured way (**Figure S6; Table S1**). For example, Col(BS512), ColKp3, IncFIA (HI1), Cml, EreA/B and GyrA 83Y/87A co-occurred frequently within the ST147-KL10 clade. Associations of (i) VIM, QnrA1, AadB and IncHI2; (ii) OXA-48-like and IncL(pOXA-48) or (iii) KPC and IncFIB(pQIL) were also observed, consistent with previous descriptions of genetic elements co-carrying these genes (Bowers et al., 2015; Coelho et al., 2012; van der Bij et al., 2011). In addition, genes conferring tolerance to copper and silver were associated with IncFII_K_/IncFIB_K_ plasmids (*pco* and *sil* operons; *p*<0.00001), whereas genes for mercuric resistance were linked to IncR plasmids (*p*=0.0082). Similarly, a positive association between the tellurite cluster and IncHIB/IncFIB (MAR) was observed (*p*<0.0001) (Navon-Venezia et al., 2017). Last, a negative association between IncR and IncFII_K_/IncFIB_K_ was observed.

### CRISPR-Cas systems

We investigated whether resistance gene dynamics could be influenced by CRISPR-Cas systems in CG147. There were either one (83%; 180/218) or two (17%; 37/218) CRISPR-Cas systems amongst CG147 genomes. A conserved subtype IV-E system was found in all genomes, located in the *iap – cysH* region, with a %G+C of 60.6%, and defined arbitrary as CRISPR1. Direct repeat sequences were highly conserved but the number and sequences of spacers varied across CG147 genomes. CRISPR1 variant v0 (43 spacers) was present in 56% of the genomes (122/218) and was used as reference to define CRISPR1 variants (**Figure S7; TableS2**). Four of the CRISPR1 v0 spacers (spacers 1, 28, 29 and 42) matched sequences on MDR IncF plasmids disseminated among different *Enterobacteriaceae*, including Kp. Spacer 1 matched a multicopy intergenic region, whereas spacers 28 and 29 matched a DUF3560-domain containing protein and spacer 42 a hypothetical protein located upstream of SAM-methyltransferase. Although these spacers were highly conserved among CRISPR1 variants, they were also found at high frequency (in 28% to 53% of the strains) as protospacers located in plasmid contigs. Noteworthy, in 28% (62/218) of CG147 genomes these plasmid protospacers were not detected, and the majority of these strains (65%, 40/62) belonged to subclades 1 and 2, which were also characterized by the absence of IncFII(K) and IncFIB(KpQIL) plasmid replicons. The association of the lack of IncF plasmids with IncF-targeting spacers in CRISPR1, suggests a possible activity of this CRISPR system in subclades 1 and 2. Finally, twelve of the forty-three spacers from CRISPR1 v0 targeted prophages; of these, spacers 6 and 25 were found as protospacers in some of the CG147 genomes (28% and 18%, respectively).

Six other CRISPR-Cas systems, distinct from CRISPR1, were observed in 37 genomes. CRISPR2 to CRISPR4, of type IV-A3, were strongly associated with IncHIB and/or IncFIB (pNDM-MAR) plasmids (*p*<0.00001), as previously described (Newire et al., 2020). Two variants of CRISPR2 (v1; n=12; 17 spacers and v3; n=5, 25 spacers) were prevalent (**Figure S7; TableS2**). CRISPR2 to CRISPR4 systems shared identical direct repeat sequences but showed a high diversity in the number and sequences of spacers. Their tendency to target IncFII_K_/IncFIB_K_ plasmids has led to suggest a role in inter-plasmid competition (Newire et al., 2020; Pinilla-Redondo et al., 2020). However, here the presence of CRISPR2 to CRISPR4 systems was not uniformly associated with an absence of IncFII_K_/IncFIB_K_ plasmids.

### Prophage elements

CG147 genomes harbored 0 to 7 prophages (mean of 4 prophages per genome; considering only the intact ones). Some prophages were frequent, including (i) ST147-VIM1phi7.1-*like* (GenBank accession no. NC_049451; *Myoviridae* family), which was present in 90% of the genomes (196/218); (ii) *Salmonella* phage 118970_sal3-*like* (GenBank accession no. NC_031940; *Myoviridae* family; 70% [152/218]); and (iii) *Enterobacteria* phage mEp237-*like* (GenBank accession no. NC_019704; *Siphoviridae* family; 56% [121/218]) (**Figure S7**).

The N15-*like* phage-plasmid (*Siphoviridae* family) we uncovered in the genomic assembly of strain DJ was present in 37% of CG147 genomes (81/218). It was strongly associated (*p*<0.00001) with ST147-KL64 subclades 1 and 2 and present in 92% of these genomes **(Table S1, Figure S7**). These two subclades were also enriched in other prophages (mean number of prophages: 5) compared with the remaining CG147 genomes (mean number of prophages=3).

## DISCUSSION

This study was triggered by the discovery of a pandrug resistance phenotype in strain DJ from India, which prompted us to analyzed its genomic features and understand their dynamics in the context of a large genome dataset of CG147 from multiple world regions. The results revealed its deep phylogenetic structure and a capacity of CG147 members to acquire a wide range of antimicrobial resistance or virulence elements, plasmids and prophages. Of these, *Klebsiella* capsular locus (KL) switches, which occurred repeatedly, represent prominent phylogenetic markers of CG147 clades. The evolutionary dynamics of these surface structure in CG147 echo those observed in other MDR clonal groups, such as ST258 or ST307 (Bowers et al., 2015; Wyres, Hawkey, et al., 2019; Wyres, Wick, et al., 2019). The multiple clonal expansions of sublineages with distinct KL or O types has clear implications for diagnostic, control or therapeutic strategies such as vaccination and phage therapy.

The genomic arsenal of antimicrobial resistance features of subclade 2, to which strain DJ belongs, suggest extensively drug and pandrug resistance is not restricted to strain DJ, but rather is a shared characteristic in this particular subclade, already disseminated among Asian countries (Alfaresi, 2018; Falcone et al., 2020; Nahid et al., 2017; Simner et al., 2018; Sonnevend et al., 2017; Zowawi et al., 2015). Clonal spread of this subclade between different countries was reported (Sonnevend et al., 2017). This subclade should be closely monitored and may represent a pioneering situation ushering the worrying prospect of pan-resistance in other CG147 subclades.

In the case of strain DJ, we observed the integration of *bla*_NDM-5_ in the chromosome. Previous studies have reported the one or multiple copies of *bla*_CTX-M-15_ and *bla*_OXA-181_ integrated into the chromosome of Kp isolates (Pitout et al., 2020; Yoon et al., 2020), including in the high-risk ST147 (Rojas et al., 2017; Sonnevend et al., 2017). However, the chromosomal integration of *bla*_NDM_ was, to our knowledge, only reported once in Kp, in two NDM-1 producing ST14 clinical isolates from Thailand (Sakamoto et al., 2018). In that case, the chromosomal integration was mediated by IS*5* and the Tn*3* transposase. In strain DJ, the integration of *bla*_NDM-5_ may have been mediated by IS*Ecp*1/IS*26* (**Figure 1a**). The genomic rearrangement due to the replicative transposition of IS*26* has previously been shown for IncFII-*bla*_NDM-5_ bearing plasmids (Takeuchi et al., 2018). The chromosomal incorporation of carbapenemase and ESBL genes is concerning, as it may stabilize these genes by promoting their vertical dissemination (Sonnevend et al., 2017).

The convergence of MDR and hypervirulence genotypes is being increasingly observed in Kp (Wyres et al., 2020). Here, this worrisome association was detected in two phylogenetically distinct CG147 isolates from 2014 (Russia) and 2016 (United Kingdom), which shared an MDR-Hv IncHIB/FIB plasmid (J. F. Turton et al., 2018). The recent observation of this plasmid in ST101 and ST147 isolates from the United Kingdom (J. Turton et al., 2019), with no epidemiological link with the isolate from 2016, suggests its continuous circulation and further risk of horizontal spread.

The ST147-KL64 lineage is globally disseminated. The evolutionary rate we estimated (1.03×10^−6^ substitutions/site/year) is very similar to other MDR global sublineages, such as ST258 (1.03×10^−6^ substitutions/site/year) and ST307 (1.18×10^−6^ substitutions/site/year) (Bowers et al., 2015; Wyres, Hawkey, et al., 2019), and slightly slower than the one estimated for ST101 (2.85×10^−6^ substitutions/site/year) (Roe et al., 2019). It is striking that the emergence of ST147-KL64 lineage occurred approximately at the same time as other MDR sublineages ST258 (year 1995), ST307 (1994) and ST101 (1989) (Bowers et al., 2015; Roe et al., 2019; Wyres, Hawkey, et al., 2019). In addition, the presence of GyrA and ParC QRDR alterations is a common characteristic to these MDR high-risk sublineages [CG258 and ST307: GyrA83I and ParC80I; ST101: GyrA83I, GyrA87G/N/A and ParC80I] (Bowers et al., 2015; Roe et al., 2019; Wyres, Hawkey, et al., 2019). This phenotypic and temporal conjunction points to common drivers and suggests a role of the usage of fluoroquinolones, introduced into clinical practice in the end of the 1980s, in the emergence of MDR Kp sublineages. This is reminiscent of the scenarios of emergence of *Escherichia coli* ST131 and methicillin-resistant *Staphylococcus aureus* ST22 (Fuzi et al., 2020; Holden et al., 2013; Stoesser et al., 2016).

The drivers of genomic diversification of emerging Kp sublineages include a combination of ecological opportunities to acquire genetic elements, local selective pressure, and molecular mechanisms that enable or restrict genetic flux. Of these, CRISPR-Cas systems may play a role (Kamruzzaman & Iredell, 2020; Li et al., 2018; Ostria-Hernández et al., 2015; Shen et al., 2017). In CG258, an association was suggested between the absence of these systems and the ability to acquire IncF plasmids [such as *bla*_KPC_-IncF(pKpQIL-like plasmids)] (Mackow et al., 2019; Tang et al., 2020; Zhou et al., 2020). Here we found a conserved type IV-E CRISPR-Cas system (CRISPR1) within CG147, with four spacers matching IncF plasmid sequences. The distribution of CRISPR1 in the broader Kp species shows a unique association with CG147, with only two exceptions (ST2746 and ST3700; based on 1001 Kp genomes representing unique STs; selected from a dataset of 4222 genomes from NCBI, November 2018; data not shown). Type IV CRISPR-Cas systems primarily target plasmids (Pinilla-Redondo et al., 2020). However, in the majority of CG147 strains, corresponding protospacers were found in plasmid sequences, advocating that the immunity provided by this CRISPR1 system might not be fully functional. In contrast, ST147-KL64 subclades 1 and 2 were largely devoid of these protospacers and did not carry IncFIA, IncFII_K_, IncFIB_K_ and IncFIB (pQil) plasmids **(Figure S7)**. This observation may suggest a possible activity of the CRIPSR1 system in these recently emerged subclades. Future experimental studies are needed to explore this hypothesis. Noteworthy, it was also among these subclades that the N15-*like* phage-plasmid was prevalent. Among bacterial phyla, this phage-plasmid family was found almost exclusively in Kp (Pfeifer et al., 2021), and we found that 58.9% of these belonged to CG147. A possible biological role of plasmid-prophages has yet to be established.

## CONCLUSIONS

The presence of pandrug-resistant and extremely-drug resistant isolates in CG147, together with the high genetic plasticity and rapid emergence dynamics of this clone, represents a clear threat to public health. CG147 is globally disseminated but shows a strong phylogenetic structure, with different clades being associated with specific genomic features and geographical distributions. These observations underline how different variants of CG147 contribute to the major public health threat posed by Kp, and call for specific surveillance and directed control strategies of this clone and its particularly concerning clades.

A possible link between the absence of IncF plasmids and the activation of CRISPR1 defense system is intriguing. This observation calls for more work on mechanistic drivers of the flux of genetic elements across MDR bacterial lineages. Precise phylogenetic mapping and understanding of the dynamics of antimicrobial resistance features are needed to guide the development of control strategies, such as CRISPR delivery or toxic conjugation systems (Bikard et al., 2014; López-Igual et al., 2019) that target specific subsets of strains within pathogenic bacterial species.

## Acknowledgments

We thank the Plateforme de Microbiologie Mutualisée (P2M) of Institut Pasteur for Illumina sequencing.

## Authors license statement

This research was funded, in whole or in part, by Institut Pasteur and by European Union’s Horizon 2020 research and innovation programme. For the purpose of open access, the authors have applied a CC-BY public copyright license to any Author Manuscript version arising from this submission.

## Ethical approval statement

To conduct the research, we used bacterial strain DJ, which is not considered a human sample. Accordingly, this research was not considered human research and is out of the scope of the decree n° 2016-1537 of November 16, 2016 implementing law n° 2012-300 of March 5, 2012 on research involving human subjects. Therefore, no ethics approval was needed and no informed consent was required. No personal identifying data was used; therefore, no consent was necessary.

## Author contributions

S.D. and D.G. coordinated the microbiological cultures of the isolate DJ and its antimicrobial susceptibility testing and biochemical characterization. V.P., C.R. and S.D. performed the genomic sequencing. C.R. designed and coordinated the comparative genomics study, with input from S.D. C.R. wrote the initial version of the manuscript. All authors provided input to the manuscript and reviewed the final version.

