## Supplementary Figures for "Genomic evolution of the globally disseminated multidrug-resistant *Klebsiella pneumoniae* clonal group 147"

**Supplementary Figure legends:**

**Figure S1**. **Phylogenetic structure of CG147.**

The tree was obtained by maximum likelihood analysis (IQ-TREE) based on the final recombination-free alignment of concatenated nucleotide sequence alignments of 4,517 core genes. The tree was rooted with a Kp ST258 NJST258_2 (GCF_000597905.1) and a Kp ST37 INF042 (GCF_002752995.1). Branch lengths represent the number of nucleotide substitutions per site (scale, 0.001 substitution per site). The three main branches are colored by sequence type (ST) and branch tips are colored by world region of isolation (see key). Capsular (KL) and O-antigen (O) locus types and the yersiniabactin-carrying ICE*Kp* elements are colored according to their variants (see key). Subclades 1 and 2 are denoted by rectangle outlines.

**Figure S2. Temporal signal in genomic sequences.**

The linear correlation between year of isolation and root-to-tip distance from the maximum likelihood phylogeny for CG147 genomes collection was calculated using TempEst.

**Figure S3**. **Replicons of strain DJ.**

(**a**) Visual representation of the hybrid Unicycler assembly graph, obtained using Bandage (Wick et al., 2015). Nine replicons were obtained, including the circularized chromosome, 6 circularized plasmids, 1 linear phage-plasmid and 1 non-circularized plasmid contig. The size of replicons is shown.

(**b**) Sequence alignments of the chromosomes of strain DJ and the four closest genomes belonging to subclade 2 (<28 SNPs): MS6671 (GenBank Accession Number: GCA_001455995), DA48896 (GCA_003006175.1); CRKP-1215 (GCA_002786755; =CRKP-

2297) and SKGH01 (GCA_001644765.1). Strain DJ was used as reference. The outermost circle is an annotation of the reference strain showing the location of antimicrobial drug resistance genes, ytb/ICEKp, KL64 capsular locus and the IncFII chromosomal integration region. The figure was created using BRIG (Alikhan et al., 2011).

(**c**) Sequence alignments of the non-circular IncFII replicon of strain DJ with the closest plasmids of subclade 2 strains: pCRKP-2297_2 (CP024836.1); pCRKP-1215_2 (CP024840.1), MS6671_plasmidE (LN824138.1), SKGH01_p3 (CP015503.1), p48896_1 (CP024430.1) and the region integrated in the chromosome of strain DJ. The IncFII non-circularized replicon was used as reference. The outermost circle is an annotation of the reference strain showing the location of antimicrobial drug resistance genes in red. In the table the coverage, identity, size and antimicrobial resistance genes of the plasmids that were used for the comparison. In bold the resistance genes also present in DJ replicon.

(**d**) Sequence alignments of the p48896_1 (CP024430.1) plasmid with the plasmids of the closest strains: pCRKP-2297_2 (CP024836.1); pCRKP-1215_2 (CP024840.1), MS6671_plasmidE (LN824138.1), SKGH01_p3 (CP015503.1), DJ-IncFII non-circularized replicon and DJ-IncR plasmid. p48896_1 was used as reference plasmid. The outermost circle is an annotation of the reference strain showing the location of antimicrobial drug resistance genes in red.

(**e**) Sequence alignments of the 123 kb IncFII(pKPX1) plasmid of strain DJ with the closest plasmids of subclade 2 strains: pCRKP-2297_1 (CP024835.1), pCRKP-1215_1 (CP024839.1), MS6671_plasmidB (LN824135.1), SKGH01_p2 (CP015502.1), p48896_2 (CP024431.1). The IncFII(pKPX1) plasmid of strain DJ was used as reference. The outermost circle is an annotation of the reference strain showing the location of antimicrobial drug resistance genes in red. In the table the coverage, identity, size and antimicrobial resistance genes of the plasmids that were used for the comparison. In bold the resistance genes also present in DJ replicon.

**Figure S4**. Geographic origins of the genomes included in this study and their associated carbapenemase profile. The pie charts represent the frequency of each carbapenemase profile in each country (see key). CPKP-: no carbapenemase.

**Figure S5.** Sequence alignments of MDR-Hv plasmids identified in CG147 genomes (KpvST147L_NDM and B-8658) and comparison with the reference virulence plasmid pK2044 (AP006726.1) and a similar plasmid recovered in 2019 (pKpvST147B, CP040726.1), which was used as reference. The outermost circle shows annotations of the reference strain.

**Figure S6**. Correlation between genotypes (Pearson method). Blank squares represent correlations without statistical significance ($p > 0.05$). The plot was created with the corrplot R package.

**Figure S7**. Time-scaled phylogeny of CG147 genomes and their heavy metal tolerance genes, plasmid replicons, CRIPSR systems, and prophages profile.

The phylogeny was obtained using the BEAST tool. The three main branches correspond to the three main sequence types (ST). Tree tips are colored by world region of isolation (see key). Black dots on main nodes indicate ≥95% posterior probability. The grey boxes delineate subclades 1 and 2, as indicated. Capsular (KL) and O-antigen (O) locus types and the yersiniabactin-carrying ICE*Kp* elements are colored according to their variants as shown in the legend. The presence of heavy metal tolerance genes, plasmid replicons, CRISPR-Cas systems, protospacers and prophages is indicated. In heavy metal tolerance genes columns, lighter yellow indicates an incomplete operon. In prophages lighter orange indicates an questionable prophage. HMTG, heavy metal tolerance genes

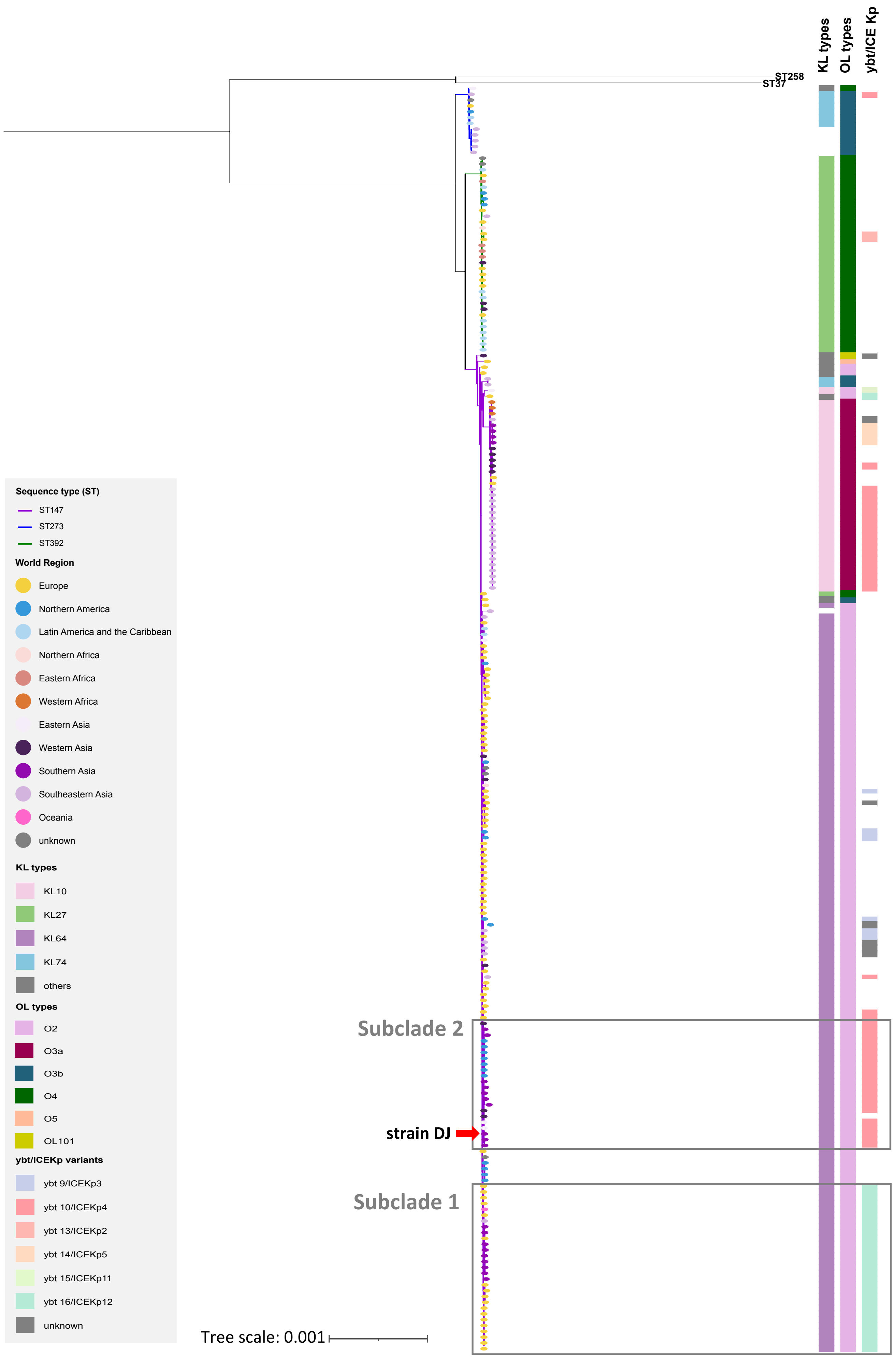
ST258
ST37
KL types
OL types
ybt/ICE Kp
Sequence type (ST)
ST147
ST273
ST392
World Region
Europe
Northern America
Latin America and the Caribbean
Northern Africa
Eastern Africa
Western Africa
Eastern Asia
Western Asia
Southern Asia
Southeastern Asia
Oceania
unknown
KL types
KL10
KL27
KL64
KL74
others
OL types
O2
O3a
O3b
O4
O5
OL101
ybt/ICEKp variants
ybt 9/ICEKp3
ybt 10/ICEKp4
ybt 13/ICEKp2
ybt 14/ICEKp5
ybt 15/ICEKp11
ybt 16/ICEKp12
unknown
Subclade 2
strain DJ
Subclade 1
Tree scale: 0.001

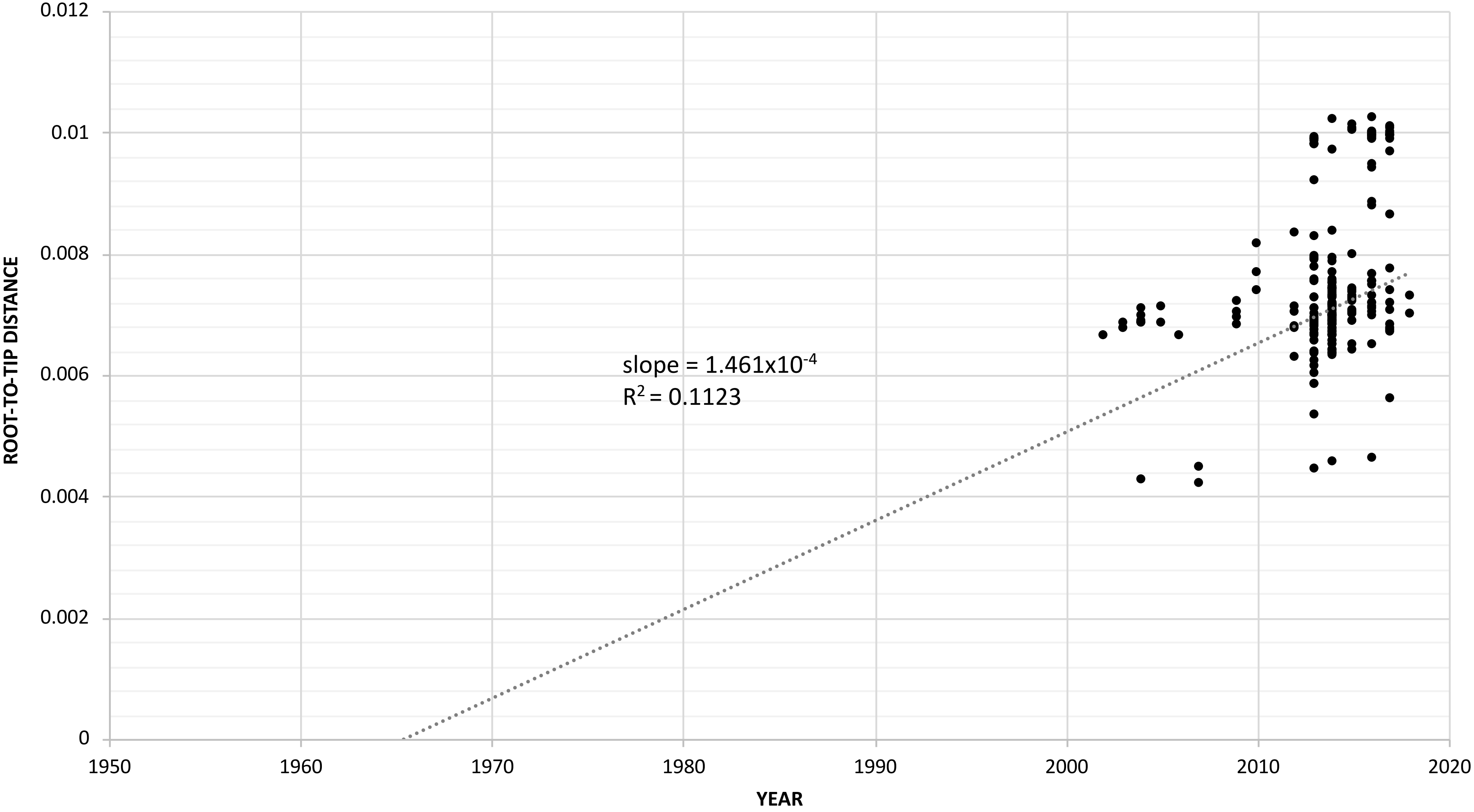

0.012
0.01
0.008
0.006
0.004
0.002
0
ROOT-TO-TIP DISTANCE
1950
1960
1970
1980
1990
2000
2010
2020
YEAR
slope = 1.461x10^-4
R^2 = 0.1123

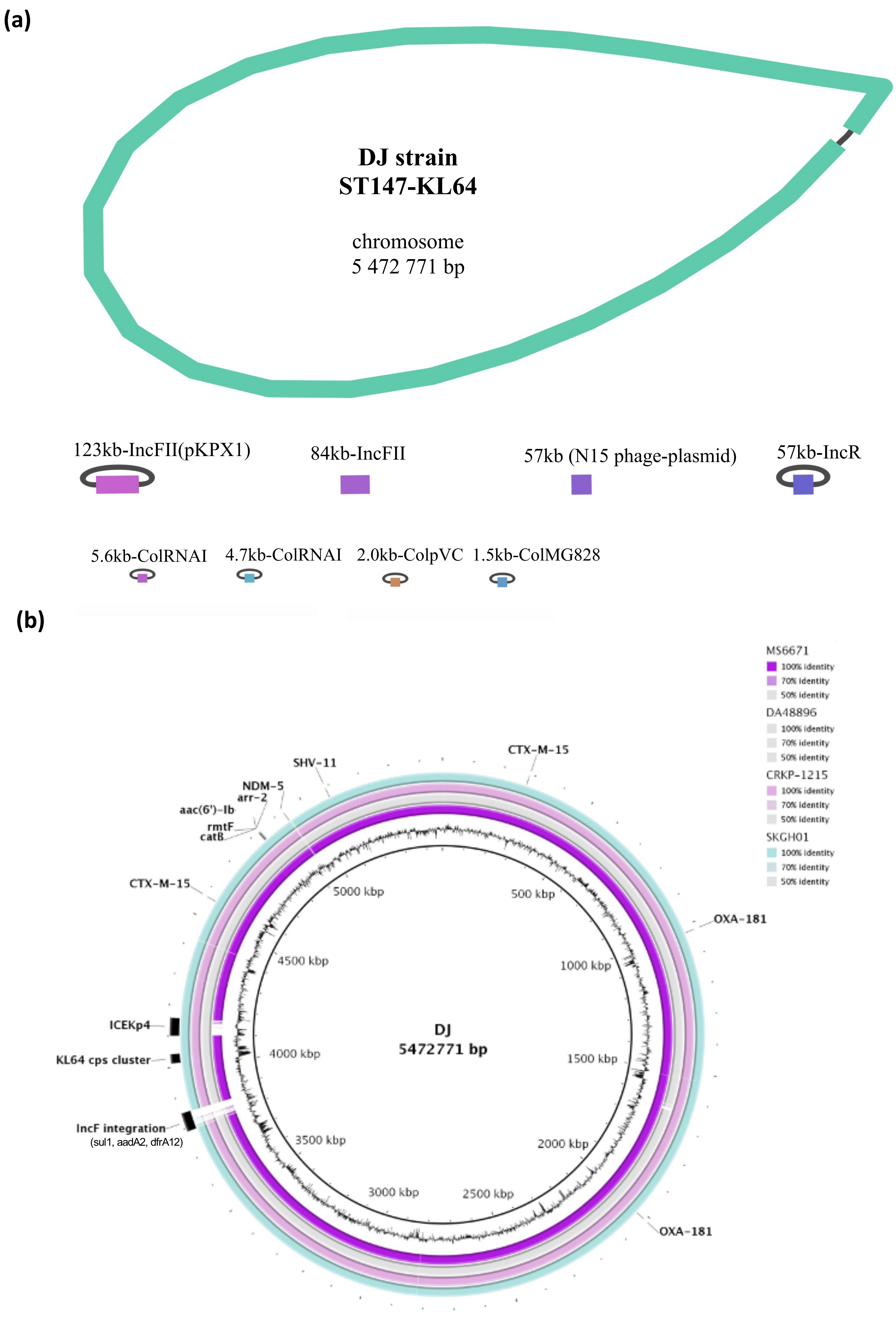
(a)
DJ strain
ST147-KL64
chromosome
5 472 771 bp
123kb-IncFII(pKPX1)
84kb-IncFII
57kb (N15 phage-plasmid)
57kb-IncR
5.6kb-ColRNAI
4.7kb-ColRNAI
2.0kb-ColpVC
1.5kb-ColMG828
(b)
MS6671
100% identity
70% identity
50% identity
DA48896
100% identity
70% identity
50% identity
CRKP-1215
100% identity
70% identity
50% identity
SKGH01
100% identity
70% identity
50% identity
CTX-M-15
SHV-11
NDM-5
arr-2
aac(6')-Ib
rmtF
catB
CTX-M-15
OXA-181
ICEKp4
KL64 cps cluster
IncF integration
(sul1, aadA2, dfrA12)
OXA-181
DJ
5472771 bp
5000 kbp
500 kbp
4500 kbp
1000 kbp
4000 kbp
1500 kbp
3500 kbp
2000 kbp
3000 kbp
2500 kbp

(c)

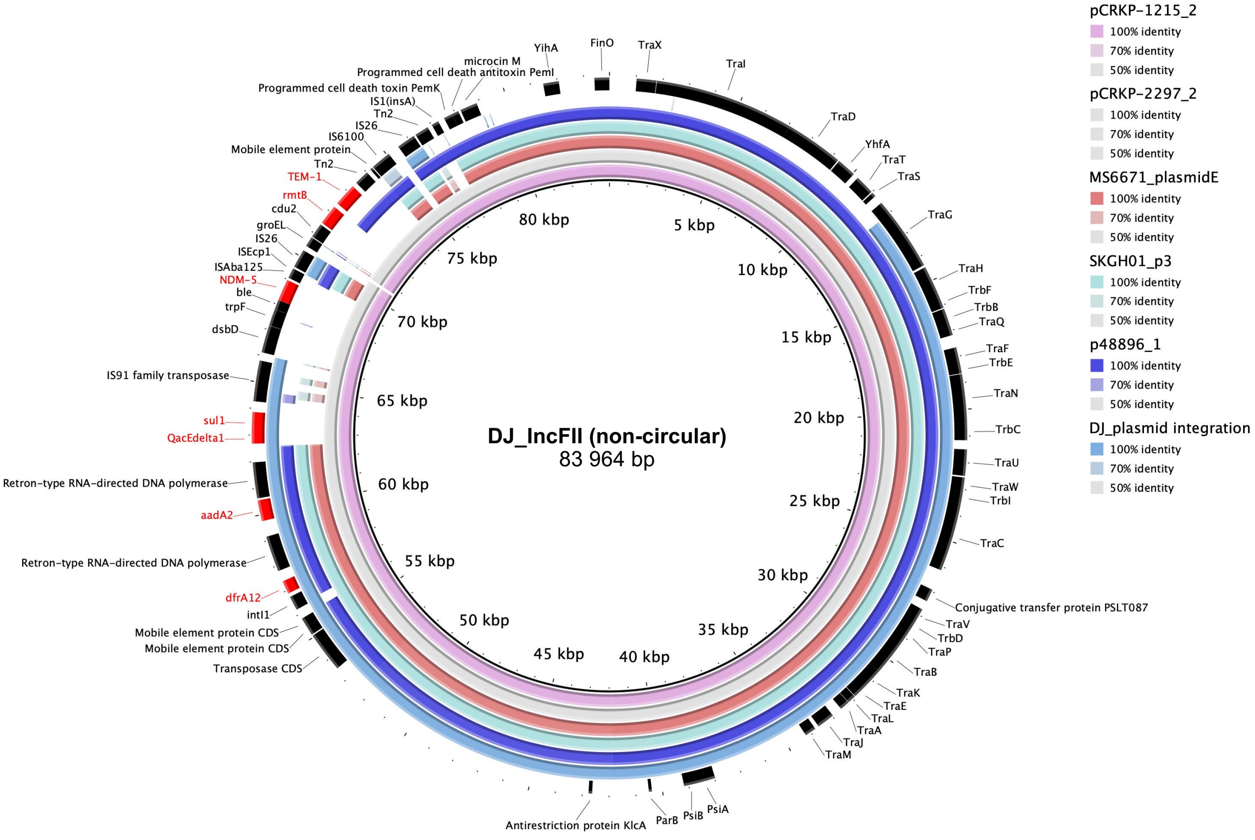


| | Cov. | ID | Size (bp) | AMR genes |
|---|---|---|---|---|
| **pCRKP-2297_2** (IncFII) | 99% | 99.61% | 96 185 | ***blaNDM−5, blaTEM−1B, dfrA12, aadA2, sul1, rmtB****, mphA, ermB* |
| **pCRKP-1215_2** (IncFII) | 99% | 99.61% | 96 185 | ***blaNDM−5, blaTEM−1B, dfrA12, aadA2, sul1, rmtB****, mphA, ermB* |
| **SKGH01_p3** (IncFII) | 85% | 99.85% | 84 941 | *aac(6')-Ib-cr; rmtF,* ***aadA2, dfrA12*** |
| **MS6671_plasmidE** (IncFII) | 85% | 99.61% | 84 940 | *aac(6')-Ib-cr; rmtF,* ***aadA2, dfrA12*** |
| **p48896_1** (IncFII/IncR) | 88% | 99.61% | 131 243 | ***blaCTX-M-15, blaTEM−1B, catA2, aadA2,*** **dfrA12, strAB (2x), sul2** |

(d)

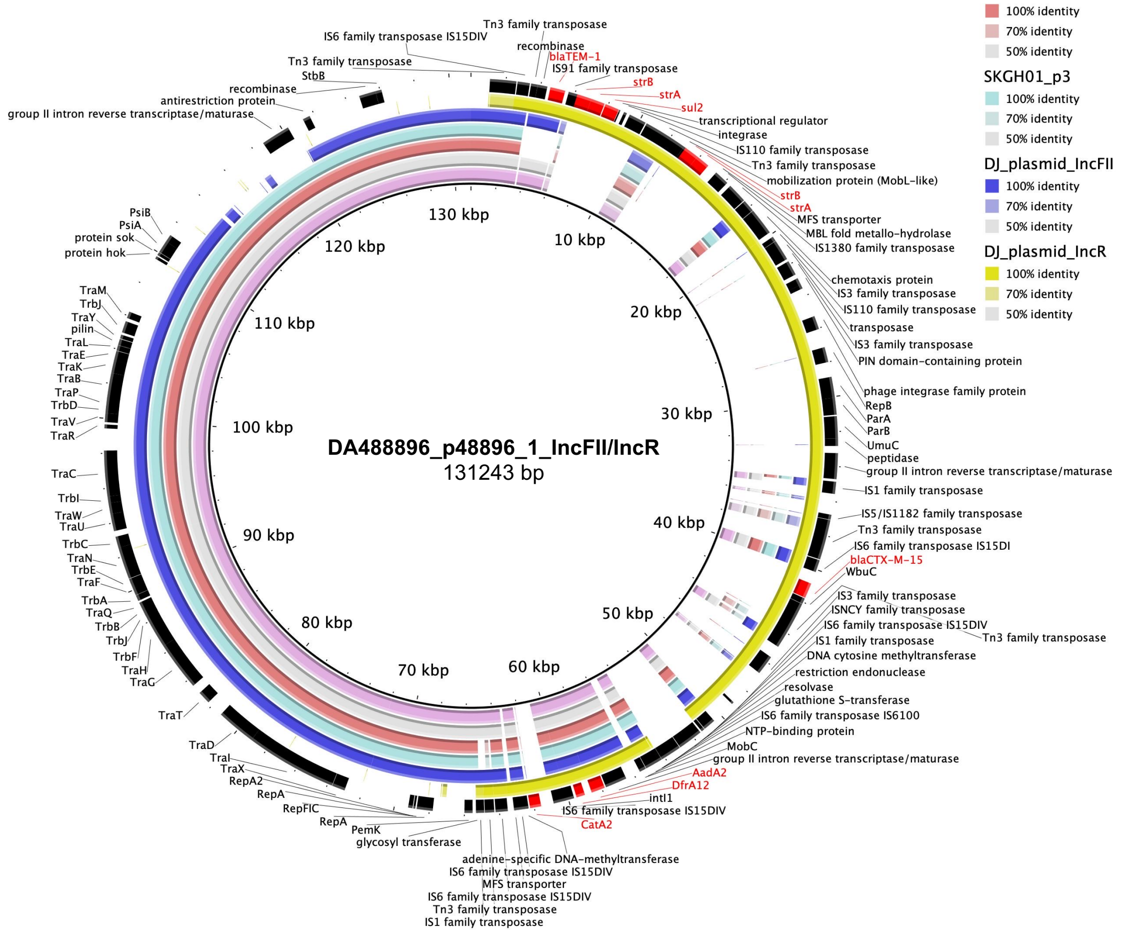

**(e)**

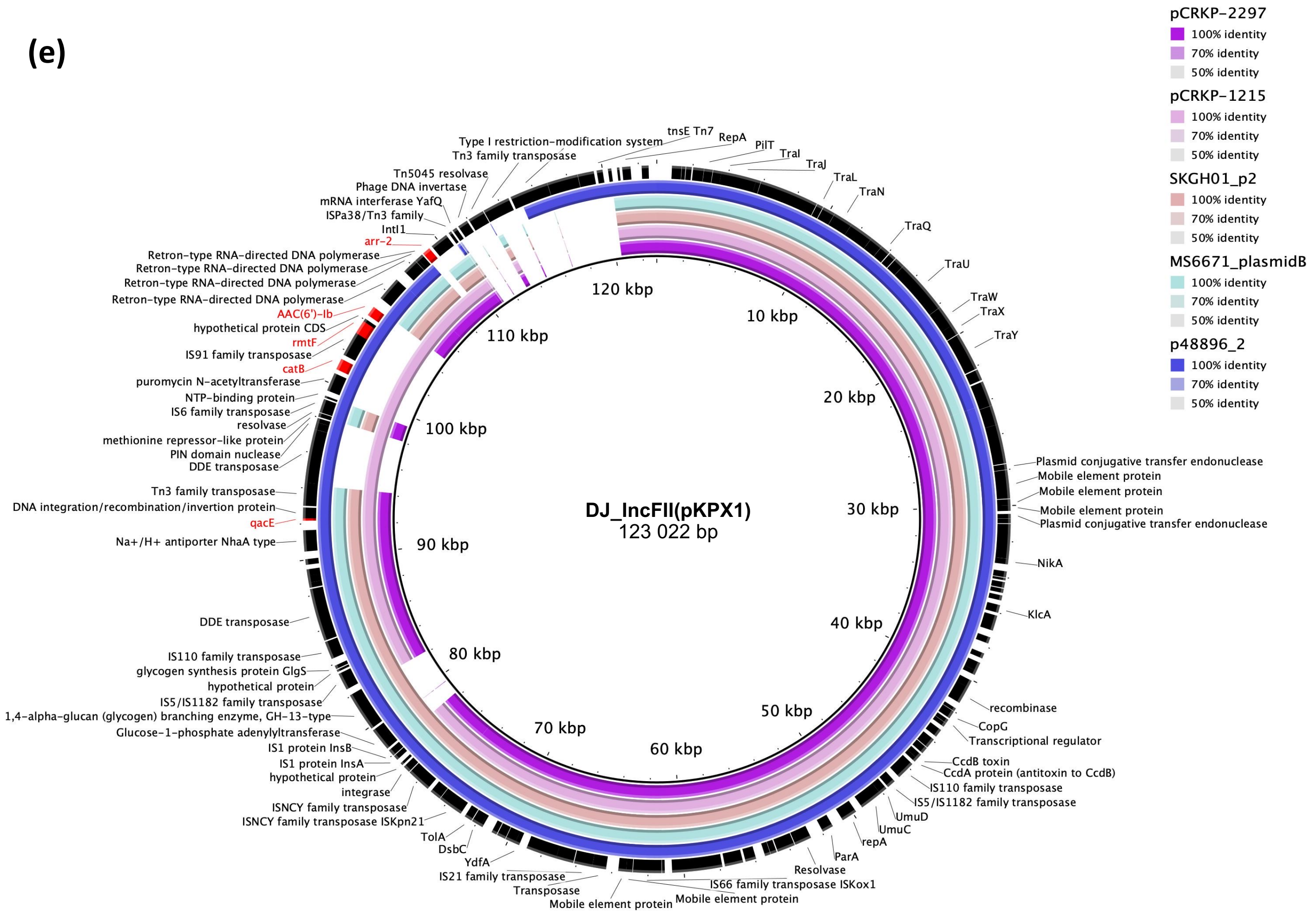


| | Cov. | ID | Size (bp) | AMR genes |
|---|---|---|---|---|
| **pCRKP-2297_1** | 81% | 99.98% | 112 150 | ***arr-2***, *qnrB1* |
| **pCRKP-1215_1** | 89% | 99.98% | 130 922 | ***arr-2***, *qnrB1*, ***rmtF***, *aadA2*, ***aacA4,*** *dfrA12*, *catA2* |
| **SKGH01_p2** | 84% | 99.78% | 140 874 | *qnrB1*, *dfrA14* |
| **MS6671_plasmidB** | 84% | 99.96% | 118 323 | *qnrB1*, *dfrA14* |
| **p48896_2** | 95% | 99.97% | 114 815 | ***rmtF, aacA4*** |

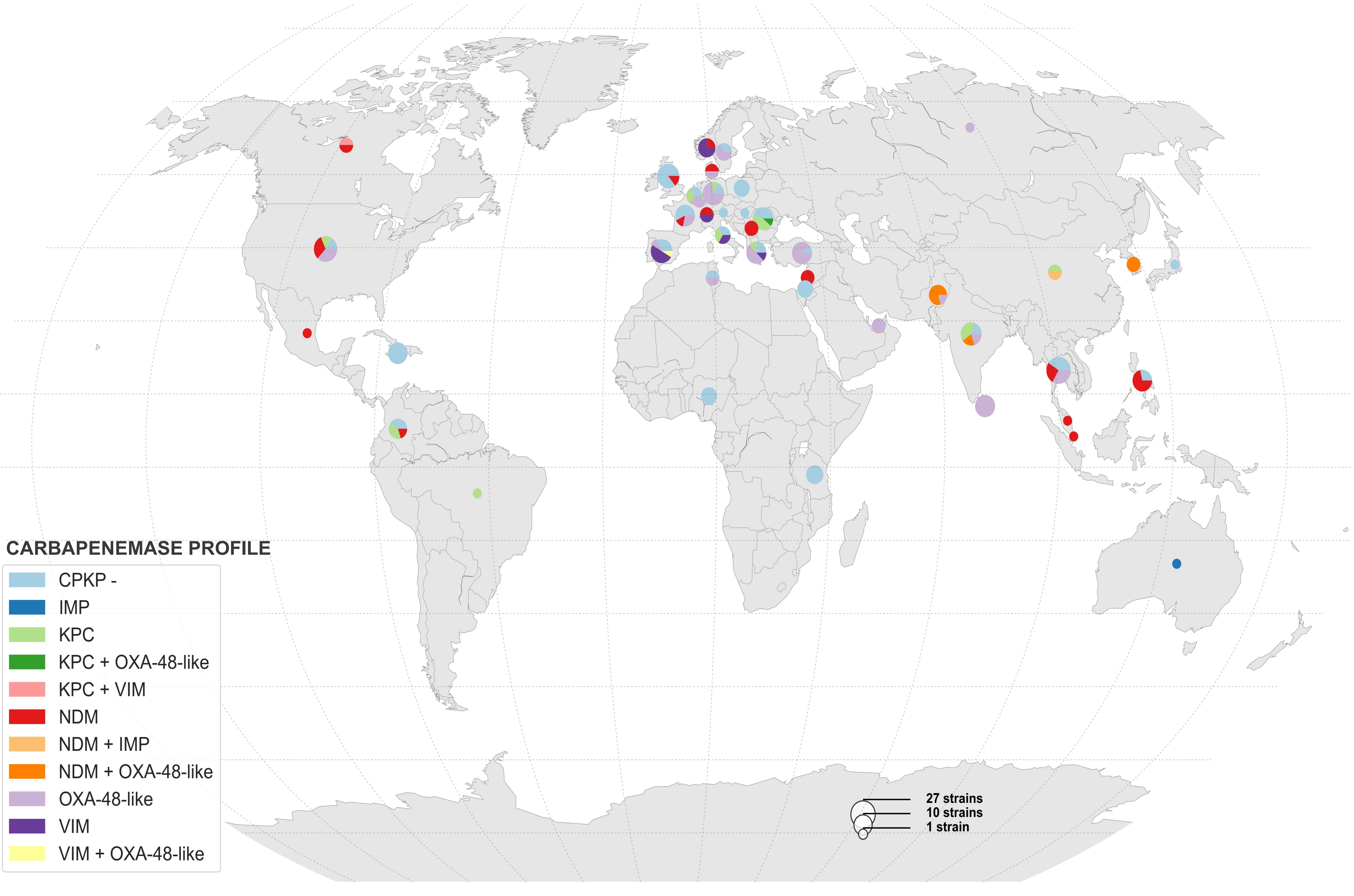
CARBAPENEMASE PROFILE
CPKP -
IMP
KPC
KPC + OXA-48-like
KPC + VIM
NDM
NDM + IMP
NDM + OXA-48-like
OXA-48-like
VIM
VIM + OXA-48-like
27 strains
10 strains
1 strain

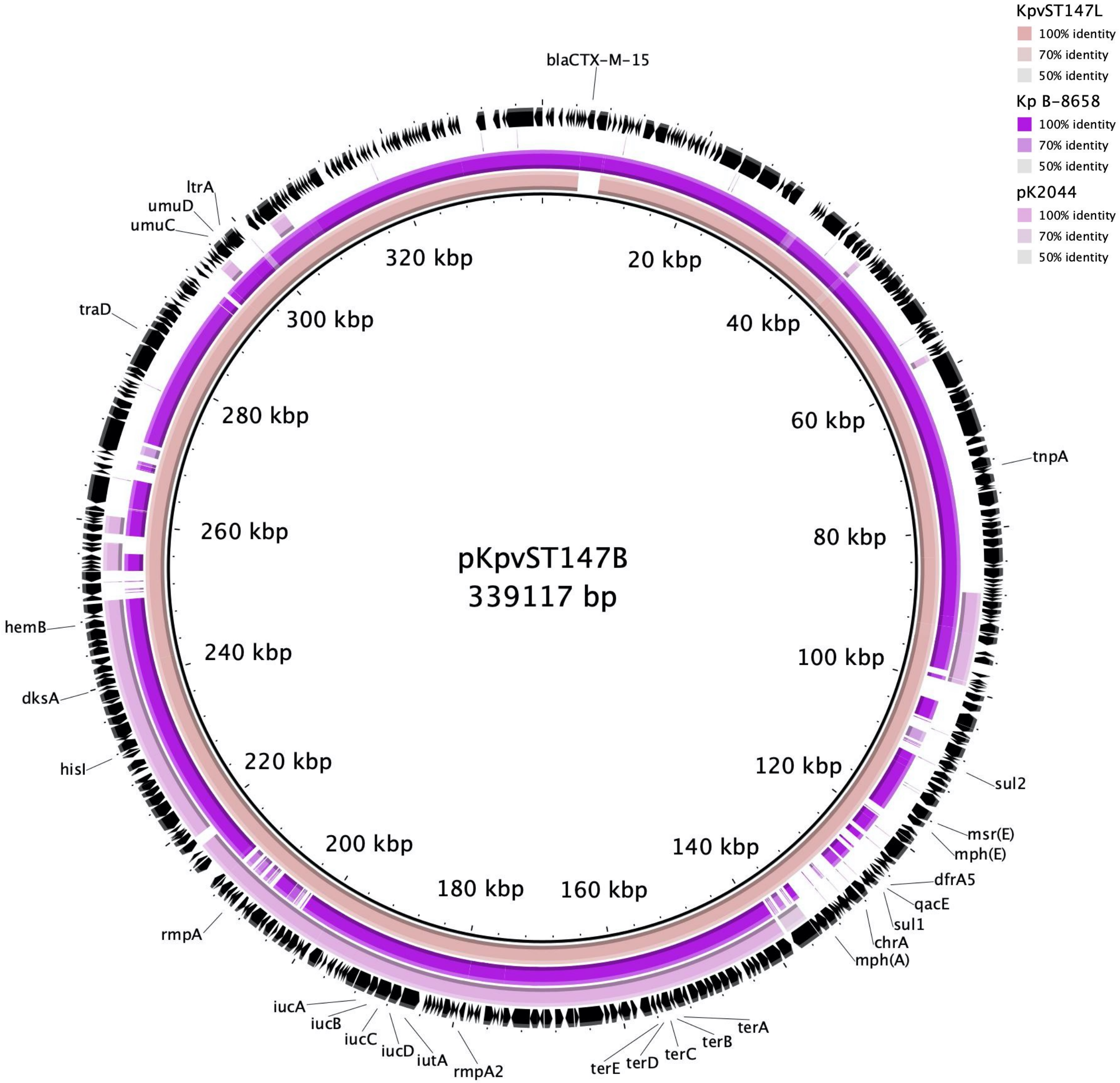

pKpvST147B
339117 bp
blaCTX-M-15
20 kbp
40 kbp
60 kbp
80 kbp
100 kbp
120 kbp
140 kbp
160 kbp
180 kbp
200 kbp
220 kbp
240 kbp
260 kbp
280 kbp
300 kbp
320 kbp
tnpA
sul2
msr(E)
mph(E)
dfrA5
qacE
sul1
chrA
mph(A)
terA
terB
terC
terD
terE
rmpA2
iutA
iucD
iucC
iucB
iucA
rmpA
hisI
dksA
hemB
traD
umuC
umuD
ltrA
KpvST147L
100% identity
70% identity
50% identity
Kp B-8658
100% identity
70% identity
50% identity
pK2044
100% identity
70% identity
50% identity

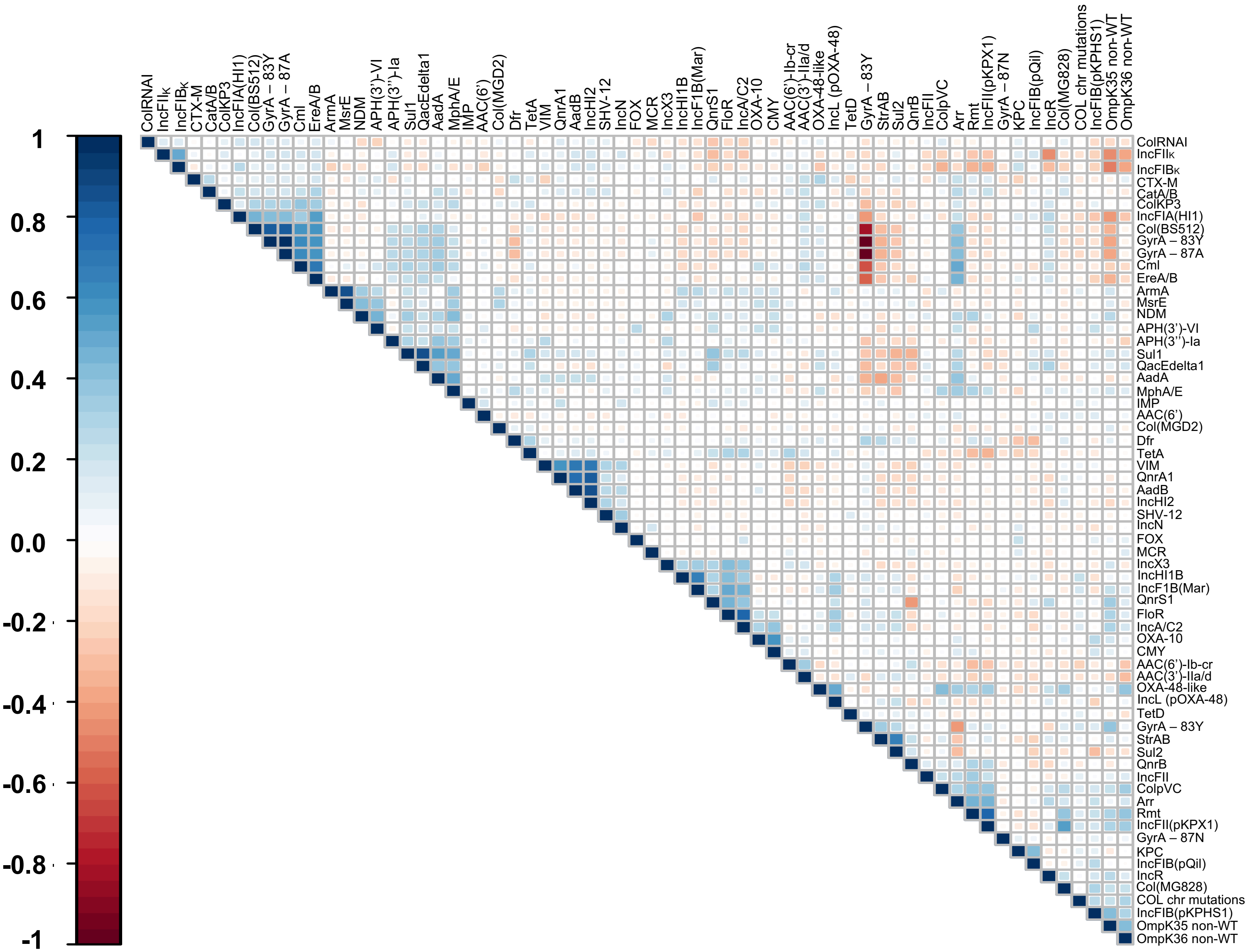

1
0.8
0.6
0.4
0.2
0.0
-0.2
-0.4
-0.6
-0.8
-1
ColRNAI
IncFIIK
IncFIBK
CTX-M
CatA/B
ColKP3
IncFIA(HI1)
Col(BS512)
GyrA – 83Y
GyrA – 87A
CmI
EreA/B
ArmA
MsrE
NDM
APH(3')-VI
APH(3'')-Ia
Sul1
QacEdelta1
AadA
MphA/E
IMP
AAC(6')
Col(MGD2)
Dfr
TetA
VIM
QnrA1
AadB
IncHI2
SHV-12
IncN
FOX
MCR
IncX3
IncHI1B
IncF1B(Mar)
QnrS1
FloR
IncA/C2
OXA-10
CMY
AAC(6')-Ib-cr
AAC(3')-IIa/d
OXA-48-like
IncL (pOXA-48)
TetD
GyrA – 83Y
StrAB
Sul2
QnrB
IncFII
ColpVC
Arr
Rmt
IncFII(pKPX1)
GyrA – 87N
KPC
IncFIB(pQil)
IncR
Col(MG828)
COL chr mutations
IncFIB(pKPHS1)
OmpK35 non-WT
OmpK36 non-WT

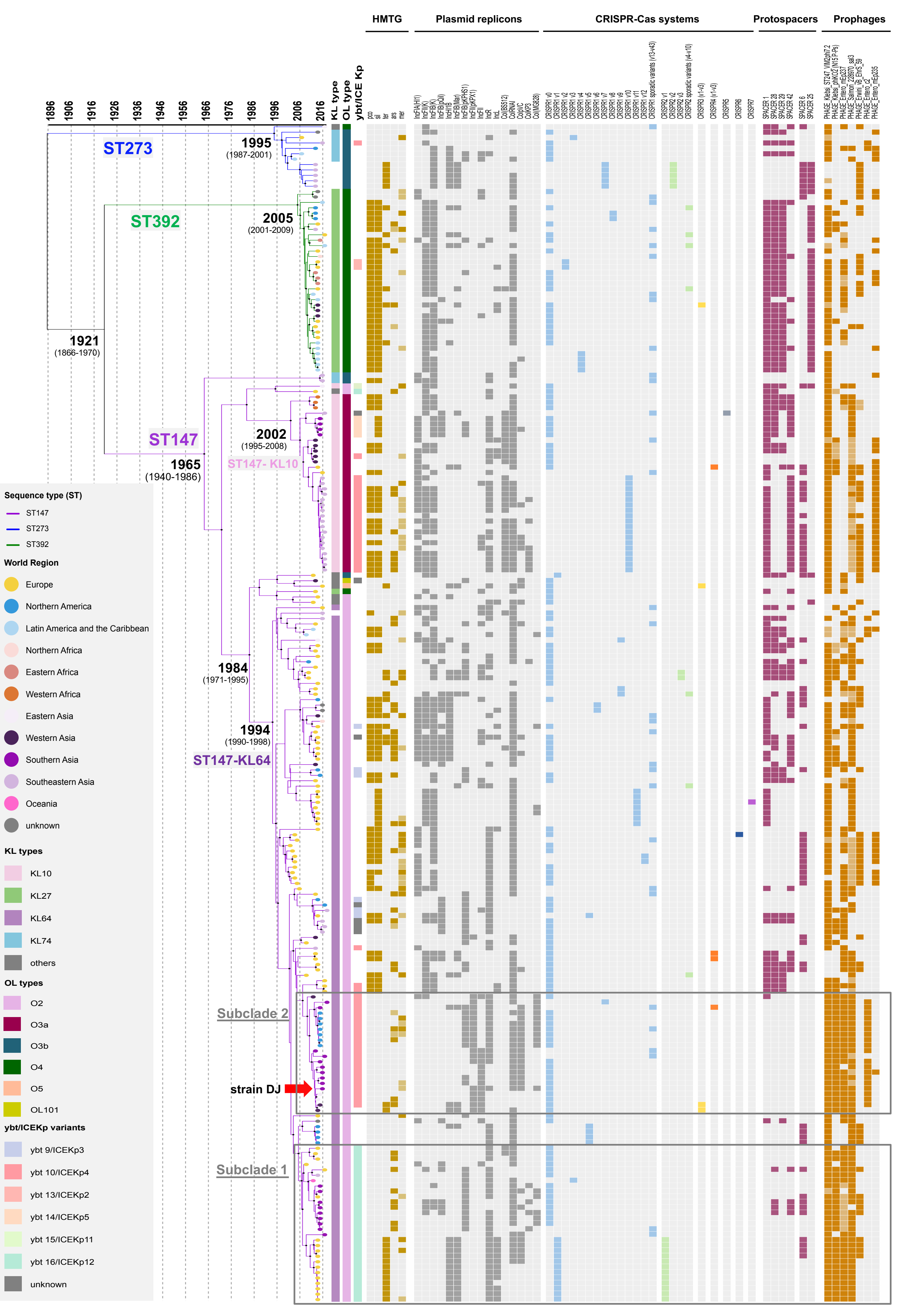

HMTG
Plasmid replicons
CRISPR-Cas systems
Protospacers
Prophages
1896
1906
1916
1926
1936
1946
1956
1966
1976
1986
1996
2006
2016
KL type
OL type
ybt/ICE Kp
ST273
1995
(1987-2001)
ST392
2005
(2001-2009)
1921
(1866-1970)
ST147
2002
(1995-2008)
1965
(1940-1986)
ST147- KL10
1984
(1971-1995)
1994
(1990-1998)
ST147-KL64
Subclade 2
strain DJ
Subclade 1
Sequence type (ST)
ST147
ST273
ST392
World Region
Europe
Northern America
Latin America and the Caribbean
Northern Africa
Eastern Africa
Western Africa
Eastern Asia
Western Asia
Southern Asia
Southeastern Asia
Oceania
unknown
KL types
KL10
KL27
KL64
KL74
others
OL types
O2
O3a
O3b
O4
O5
OL101
ybt/ICEKp variants
ybt 9/ICEKp3
ybt 10/ICEKp4
ybt 13/ICEKp2
ybt 14/ICEKp5
ybt 15/ICEKp11
ybt 16/ICEKp12
unknown
