## Supplementary material for "Genomic evolution of the globally disseminated multidrug-resistant *Klebsiella pneumoniae* clonal group 147": Table S2

| **CRISPR system/variant**  **Table S2**. CRISPR/Cas systems and variants identified within CG147 *K. pneumoniae* genomes. | **No. of genomes** | **Size (bp)** | **DR consensus** | **No. of spacers** | **Comments** |
| --- | --- | --- | --- | --- | --- |
| **CRISPR1 v0** | 122 | 2650 | GTGTTCCCCGCGCCAGCGGGGATAAACCG | 43 | considered as reference |
| **CRISPR1 v1** | 13 | 2162 | GTGTTCCCCGCGCCAGCGGGGATAAACCG | 35 | missing spacers 11 to 18 |
| **CRISPR1 v2** | 2 | 2162 | GTGTTCCCCGCGCCAGCGGGGATAAACCG | 35 | missing spacers 35 to 43 |
| **CRISPR1 v3** | 2 | 2528 | GTGTTCCCCGCGCCAGCGGGGATAAACCG | 41 | missing spacers 2 and 3 |
| **CRISPR1 v4** | 4 | 2675 | GTGTTCCCCGCGCCAGCGGGGATAAACCG | 43 | duplication in one DR |
| **CRISPR1 v5** | 5 | 2589 | GTGTTCCCCGCGCCAGCGGGGATAAACCG | 42 | missing spacer 3 |
| **CRISPR1 v6** | 2 | 2223 | GTGTTCCCCGCGCCAGCGGGGATAAACCG | 36 | missing spacers 21 to 27 |
| **CRISPR1 v7** | 6 | 2406 | GTGTTCCCCGCGCCAGCGGGGATAAACCG | 39 | missing spacers 21 to 24 |
| **CRISPR1 v8** | 2 | 2406 | GTGTTCCCCGCGCCAGCGGGGATAAACCG | 39 | missing spacers 36 to 39 |
| **CRISPR1 v9** | 3 | 2528 | GTGTTCCCCGCGCCAGCGGGGATAAACCG | 41 | missing spacers 38 and 39 |
| **CRISPR1 v10** | 18 | 1735 | GTGTTCCCCGCGCCAGCGGGGATAAACCG | 28 | missing spacers 12 to 16; 21 to 29 and 38 |
| **CRISPR1 v11** | 7 | 1308 | GTGTTCCCCGCGCCAGCGGGGATAAACCG | 21 | missing spacers 17 to 38 |
| **CRISPR1 v12** | 2 | 1369 | GTGTTCCCCGCGCCAGCGGGGATAAACCG | 22 | missing spacers 9 to 29 |
| **CRISPR1 v13** | 1 | 2772 | GTGTTCCCCGCGCCAGCGGGGATAAACCG | 45 | two extra spacers; spacer 26 different |
| **CRISPR1 v14** | 1 | 2679 | GTGTTCCCCGCGCCAGCGGGGATAAACCG | 43 | missing spacer 27; spacer 26 different; extra spacer |
| **CRISPR1 v15** | 1 | 2650 | GTGTTCCCCGCGCCAGCGGGGATAAACCG | 43 | missing spacer 38; extra spacer in the beginning |
| **CRISPR1 v16** | 1 | 2616 | GTGTTCCCCGCGCCAGCGGGGATAAACCG | 42 | missing spacer 3 |
| **CRISPR1 v17** | 1 | 2589 | GTGTTCCCCGCGCCAGCGGGGATAAACCG | 42 | missing spacer 34 |
| **CRISPR1 v18** | 1 | 2589 | GTGTTCCCCGCGCCAGCGGGGATAAACCG | 42 | missing spacer 25 |
| **CRISPR1 v19** | 1 | 2589 | GTGTTCCCCGCGCCAGCGGGGATAAACCG | 42 | missing spacer 38 |
| **CRISPR1 v20** | 1 | 2558 | GTGTTCCCCGCGCCAGCGGGGATAAACCG | 41 | missing spacers 5 and 6 |
| **CRISPR1 v21** | 1 | 2553 | GTGTTCCCCGCGCCAGCGGGGATAAACCG | 41 | missing spacers 41 and 42: duplication in one DR |
| **CRISPR1 v22** | 1 | 2528 | GTGTTCCCCGCGCCAGCGGGGATAAACCG | 41 | missing spacers 26 and 27 |
| **CRISPR system/variant** | **No. of genomes** | **Size (bp)** | **DR consensus** | **No. of spacers** | **Comments** |
| **CRISPR1 v23** | 1 | 2528 | GTGTTCCCCGCGCCAGCGGGGATAAACCG | 41 | missing spacers 41 and 42 |
| **CRISPR1 v24** | 1 | 2528 | GTGTTCCCCGCGCCAGCGGGGATAAACCG | 41 | missing spacers 30 to 31 |
| **CRISPR1 v25** | 1 | 2528 | GTGTTCCCCGCGCCAGCGGGGATAAACCG | 41 | missing spacers 11 to 12 |
| **CRISPR1 v26** | 1 | 2467 | GTGTTCCCCGCGCCAGCGGGGATAAACCG | 40 | missing spacers 40 to 42 |
| **CRISPR1 v27** | 1 | 2467 | GTGTTCCCCGCGCCAGCGGGGATAAACCG | 40 | missing spacers 26 to 28 |
| **CRISPR1 v28** | 1 | 2406 | GTGTTCCCCGCGCCAGCGGGGATAAACCG | 39 | missing spacers 26 to 29 |
| **CRISPR1 v29** | 1 | 2406 | GTGTTCCCCGCGCCAGCGGGGATAAACCG | 39 | missing spacers 24 to 27 |
| **CRISPR1 v30** | 1 | 2406 | GTGTTCCCCGCGCCAGCGGGGATAAACCG | 39 | missing spacers 14-16 and 27 |
| **CRISPR1 v31** | 1 | 2344 | GTGTTCCCCGCGCCAGCGGGGATAAACCG | 38 | missing spacers 1-5 |
| **CRISPR1 v32** | 1 | 2284 | GTGTTCCCCGCGCCAGCGGGGATAAACCG | 38 | missing spacers 37 to 42 |
| **CRISPR1 v33** | 1 | 2223 | GTGTTCCCCGCGCCAGCGGGGATAAACCG | 36 | missing spacers 17; 35 to 40 |
| **CRISPR1 v34** | 1 | 2187 | GTGTTCCCCGCGCCAGCGGGGATAAACCG | 35 | missing spacers 8 and 9; 11 to 22 |
| **CRISPR1 v35** | 1 | 2100 | GTGTTCCCCGCGCCAGCGGGGATAAACCG | 34 | missing spacers 21 to 29 |
| **CRISPR1 v36** | 1 | 2040 | GTGTTCCCCGCGCCAGCGGGGATAAACCG | 33 | missing spacers 17; 31 to 33, 35 to 40 |
| **CRISPR1 v37** | 1 | 1887 | GTGTTCCCCGCGCCAGCGGGGATAAACCG | 30 | missing spacers 20 to 32 |
| **CRISPR1 v38** | 1 | 1673 | GTGTTCCCCGCGCCAGCGGGGATAAACCG | 27 | missing spacers 4, 5 and 18 to 31 |
| **CRISPR1 v39** | 1 | 1674 | GTGTTCCCCGCGCCAGCGGGGATAAACCG | 27 | missing spacers 8 to 22, and 25 |
| **CRISPR1 v40** | 1 | 1399 | GTGTTCCCCGCGCCAGCGGGGATAAACCG | 22 | missing spacers 13 to 33 |
| **CRISPR1 v41** | 1 | 1276 | GTGTTCCCCGCGCCAGCGGGGATAAACCG | 21 | missing spacers 15 to 37 |
| **CRISPR1 v42** | 1 | 1003 | GTGTTCCCCGCGCCAGCGGGGATAAACCG | 16 | missing spacers 8 to 14; and 16 to 35 |
| **CRISPR1 v43** | 1 | 575 | GTGTTCCCCGCGCCAGCGGGGATAAACCG | 9 | missing spacers 4 to 20; 23 to 34 and 37 to 41 |
| **CRISPR2 v1** | 12 | 1056 | CCGATAACCCCCGCATGCGGGGGGAATAC | 17 | - |
| **CRISPR2 v2** | 5 | 1549 | CCGATAACCCCCGCATGCGGGGGGAATAC | 25 | - |
| **CRISPR system/variant** | **No. of genomes** | **Size (bp)** | **DR consensus** | **No. of spacers** | **Comments** |
| **CRIPSR2 v3** | 2 | 515 | CCGATAACCCCCGCATGCGGGGGGAATAC | 8 | - |
| **CRISPR2 v4** | 1 | 1367 | CCGATAACCCCCGCATGCGGGGGGAATAC | 22 | - |
| **CRISPR2 v5** | 1 | 1245 | CCGATAACCCCCGCATGCGGGGGGAATAC | 20 | - |
| **CRISPR2 v6** | 1 | 939 | CCGATAACCCCCGCATGCGGGGGGAATAC | 15 | - |
| **CRISPR2 v7** | 1 | 814 | CCGATAACCCCCGCATGCGGGGGGAATAC | 13 | - |
| **CRISPR2 v8** | 1 | 761 | CCGATAACCCCCGCATGCGGGGGGAATAC | 12 | - |
| **CRISPR2 v9** | 1 | 696 | CCGATAACCCCCGCATGCGGGGGGAATAC | 11 | - |
| **CRISPR2 v10** | 1 | 456 | CCGATAACCCCCGCATGCGGGGGGAATAC | 7 | - |
| **CRISPR3 v1** | 2 | 1060 | CCGATAACCCCCGCACACGGGGGGAATAC | 17 | - |
| **CRISPR3 v2** | 1 | 1004 | CCGATAACCCCCGCACACGGGGGGAATAC | 16 | - |
| **CRISPR3 v3** | 1 | 638 | CCGATAACCCCCGCACACGGGGGGAATAC | 10 | - |
| **CRISPR4 v1** | 2 | 1238 | CCGATAACCCCCGCATGCGGGGG | 19 | - |
| **CRISPR4 v2** | 1 | 815 | CCGATAACCCCCGCATGCGGGGG | 13 | - |
| **CRISPR4 v3** | 1 | 631 | CCGATAACCCCCGCATGCGGGGG | 10 | - |
| **CRISPR5** | 1 | 1788 | CCGATAACCCCCGCAAGCGGGGGAATAC | 29 | - |
| **CRISPR6** | 1 | 599 | CCGCGCCGGTGGACGCCCCGGCCGCCGAACCGGTCGACCCGCGCAAGGCGGCGG | 5 | - |
| **CRISPR7** | 1 | 399 | AGAAACACCCCCACGTGCGTGGGGAAGAC | 5 | - |
